# Early age-related atrophy of cutaneous lymph nodes precipitates an early functional decline in skin immunity in mice with aging

**DOI:** 10.1101/2021.11.22.469479

**Authors:** Sandip Ashok Sonar, Jennifer L. Uhrlaub, Christopher P. Coplen, Gregory D. Sempowski, Jarrod A. Dudakov, Marcel R.M. van den Brink, Bonnie J. LaFleur, Mladen Jergovic, Janko Nikolich-Žugich

## Abstract

Secondary lymphoid organs (SLO; including the spleen and lymph nodes) are critical both for the maintenance of naïve T (T_N_) lymphocytes and for the initiation and coordination of immune responses. How they age, including the exact timing, extent, physiological relevance, and the nature of age-related changes, remains incompletely understood. We used time-stamping to indelibly mark cohorts of newly generated naïve T cells (a.k.a. recent thymic emigrants - RTE) in mice, and followed their presence, phenotype and retention in SLO. We found that SLO involute asynchronously. Skin-draining lymph nodes (LN) atrophied early (6-9 months) in life and deeper tissue-draining LN and the spleen late (18-20 months), as measured by the loss of both T_N_ numbers and the fibroblastic reticular cell (FRC) network. Time-stamped RTE cohorts of all ages entered SLO and successfully completed post-thymic differentiation. However, in older mice, these cells were poorly retained, and those found in SLO exhibited an emigration phenotype (CCR7^lo^S1P1^hi^). Transfers of adult RTE into recipients of different ages formally demonstrated that the defect segregates with the age of the SLO microenvironment and not with the age of T cells. Finally, upon intradermal immunization, RTE generated in mice as early as 6-7 months of age barely participated in *de novo* immune responses and failed to produce well-armed effector cells. These results highlight changes in structure and function of superficial secondary lymphoid organs in laboratory mice that are earlier than expected and are consistent with the long-appreciated and pronounced reduction of cutaneous immunity with aging.

## INTRODUCTION

Naive T (T_N_) cells are critically important for mounting *de novo* immune responses against emerging and reemerging pathogens, due to their broad TCR repertoire diversity and consequent specificity for a wide array of antigens. Ontogenically, T_N_ cells are produced and released by the thymus, and are then transported by blood and lymph to seed secondary lymphoid organs (SLO; in this manuscript, spleen and lymph nodes - LN). Such cells are called recent thymic emigrants (RTE)^1^, a definition usually applied to the cells in the first 7-14 days following thymic egress. RTE have characteristic phenotypic and functional features that make them distinct from T_N_ cells, including high threshold for TCR activation, high expression of anergy-related genes, and lower cytokine production capability upon stimulation as compared to mature T_N_ cells^2^. However, RTE are known to produce comparable effector response to that of fully mature T_N_ cells under inflammatory conditions and upon infectious challenge in mice^3^. A series of seminal studies led to a model where upon successful exit from the thymus, RTE migrate to the SLO and interact with both stromal and migrating myeloid cells (e.g. dendritic cells, DC) that provide further signals leading to their final maturation into T_N_ cells^2, 4^. It simultaneously became clear that SLO are the key sites that maintain the T_N_ cell pool under homeostatic and other conditions, including inflammation, infection and lymphodepletion^5, 6, 7, 8^. Therefore, homeostatic maintenance of T_N_ cell pool largely depends on the balance between seeding and retention of RTE, long-term maintenance of the new and existing T_N_ cells in the SLO, and their loss due to phenotypic transition into other subsets as a consequence of antigenic stimulation, altered maintenance, or death. This ensures that T_N_ cell numbers, diversity, and function are maintained for long time periods in the SLO, especially after thymic involution^9, 10^. However, homeostatic maintenance of T_N_ cells deteriorates with advanced age, potentially pronouncing the impairment of protective immunity in older adults^11, 12^.

Stromal cells of the SLO consist of many different cell types of non-hematopoietic origin which not only provide the overall structural scaffolding of the SLO, but more importantly organize themselves into a number of specialized microenvironments (or niches) that play critical roles in lymphocyte maintenance and function^13^. For example, fibroblastic reticular cells (FRC) form a network of reticular conduits coated by extracellular matrix (ECM), that provide homeostatic signals to T_N_ cells in the form of IL-7 deposited on the ECM and serve as “superhighways” to direct incoming DC and T_N_ cells to one another during the initiation of an immune response. Recent studies from some^5, 14^ but not other^15^ groups have suggested that age-related structural deterioration in SLO might hamper the maintenance and function of T_N_ cells in old mice^5, 14, 5, 16^. Specifically, Becklund et al ^5^ showed that in the steady, homeostatic state, the superficial LN of the old recipient mice exhibited a defect in the maintenance of young donor CD4^+^ and CD8^+^ T cells upon adoptive transfers. This correlated to a numerical reduction in FRC and reduced connectivity of the FRC network in the T cell zone, which could potentially reduce the bioavailability/access to survival signals, such as IL-7, as well as to a loss in demarcation between T- and B-cell zones in the LN^5^. These results were corroborated and extended by our group to describe an even more pronounced age-related numerical reduction in lymphatic endothelial cells (LEC)^13^. A similar deterioration in T- and B-cell zone organization and altered T/B interactions have also been reported in human lymph nodes^17^. With regard to initiation of new immune responses, old superficial LN fail to expand in response to West Nile virus (WNV)^16^ and Chikungunya virus (CHIKV)^18^ infections, resulting in reduced T and B cell accumulation, fewer germinal centers, and reduced levels of neutralizing antibodies^16, 18^. In the case of WNV, Richner et al. showed that immigrant T and B cells exhibit an age-related delay in entry as well as slow and apparently disorganized migration within the LN^16^, consistent with the disorganization of the FRC network architecture. On the other hand, Masters et al.^15^ reported data that confirmed some, but not all, of these results. Specifically, they found age-related changes that affected the architecture and organization of the LN, erosion of splenic gp38/podoplanin^+^ (a FRC marker) area and reduction in T cell zone FRC (also called T-cell zone reticular cells, or TRC) numbers as well as reduced levels of CCL19 and CCL21 per mg of total protein^15, 19^. However, they also reported that at a steady-state, popliteal LN (superficial) and lung-draining (deep) mediastinal LNs of the young and old mice had similar numbers of stromal cells, but that upon influenza infection the proliferation and expansion of the major stromal cell subsets (FRC (CD45^-^gp38^+^CD31^-^), LEC (CD45^-^gp38^+^CD31^+^) and BEC (CD45^-^gp38^-^CD31^+^) of the old mediastinal LN appeared delayed compared to young couterparts^15^. While these results collectively indicate some level of SLO dysfunction with aging, whether and to what extent these age-related LN structural and stromal changes contribute to defects in the maintenance and function of T_N_ remains incompletely understood.

The pool of T_N_ cells is continuously maintained by balancing influx of newly generated RTE and the homeostatic regulation of the mature T_N_ compartment in SLO. A cross-sectional analysis of RTE using Rag-2-pGFP mice revealed that GFP^+^ RTE in the spleen started declining around 24-25 weeks of age and remained stable thereafter^20^. However, in that model thymocytes were labeled by GFP expressed off the episomal circles generated during V-D-J recombination, and the label is relatively rapidly lost following exit from the thymus (typically within a week of egress). This disallows studies of cohorts of T cells born at different ages or their prolonged follow-up in the post-thymic period in SLO. Because residual thymic output remains even at older ages, it is important to understand the biology of the RTE generated *de novo* in old age and to compare them to those generated earlier in life. For these reasons, the ability of newly generated RTE to seed, survive, and be maintained in the pLN at different ages has not been systematically investigated. Zhang et. al. elegantly solved the technical limitations of this problem, by stably marking distinct waves of T cells using the tamoxifen-driven TCRδCre.ER-ZsGreen inducible labeling^21^. They followed T cells generated at early (1 mo) and late (16 mo) life and found that labeled T cells in SLO were mainly of naïve and memory phenotype, at 3 and 18 mo of age, respectively^21^, consistent with other work in the field (rev. in^11^).

To further address the timing, the extent, and physiological importance of age-related SLO defects, including disturbances in SLO stroma, we deployed the above TCRdCre.ER mouse model^21^ (generously provided by Dr Y. Zhuang, Duke University), which allowed us to “time-stamp”-label distinct waves of newly generated RTE at different chronological ages. We report a surprisingly early decline (6-9 mo) of RTE in the skin-draining (axillary and inguinal) LN, whereas the deeper SLO (e.g. brachial LN and spleen) exhibited a late-life decline. These kinetics of RTE decline corresponded closely to the reduction of LN-homing CCR7^+^ cells and with a transient increase in CCR7^-^S1P1^+^ LN emigrants, but was independent of the decline in export of RTE from the thymus. The RTE decline in different LN temporally coincided with a numerical reduction of LN LECs, as well as with a profound disruption of the LN FRC network –early in superficial LN and late in deep-draining LN and spleen. Moreover, T cells generated early in life declined numerically in SLO around 5-6 mo and acquired a virtual memory phenotype. Adoptive transfers of adult T_N_ cells into old or adult recipients allowed us to map reduced retention of RTE by LN at different ages to the age of LN and not to the intrinsic age of the T_N_ cells. Finally, functional experiments demonstrated that already at 6 months, RTE participated poorly in primary immune responses, particularly at the level of fully differentiated and armed (Granzyme B+, Ag-specific) CD8 T cells. Collectively, this data suggests that LNs do not age equally. Skin-draining LN exhibited early atrophy and deeper LN (and spleen) maintained their structure longer, providing a useful temporal framework to dissect the cellular and molecular basis for defects in T_N_ cell maintenance and function.

## RESULTS

### Differential kinetics of seeding and retention of newly generated recent thymic emigrants in the SLO

Several studies in both humans and mice have shown the decline in T_N_ cell numbers during aging ^22, 23, 24^. Using the Rag2-pGFP reporter mouse model we have shown that in the last third of life, even when the thymus is reactivated, RTE failed to home to and/or be retained well in the old SLO, particularly the pLN^14^. This may be due to defects in either the old T-cells, the pLN microenvironment, or both. Two main experimental approaches have been used to label recent thymic emigrants and study their entry and retention into SLO – direct fluorescein injection into the thymus^25^ and genetic marking of recently completed V(D)J recombination by GFP driven by the Rag2 promoter^26^. Both are very efficient in short-term labeling of RTE, however, the label disappears within days due to cell division and protein turnover, and therefore the RTE cells, defined by the Rag2-pGFP expression can be followed, at most, for up to days 4-7 post thymic exit and those labeled by fluorescein for even shorter periods.

To stably label and monitor the persistence of RTE for long periods of time, we used tamoxifen-inducible TCRδ^CreER^.Rosa26-ZsGreen reporter mice^21^. These animals were fed tamoxifen-containing diet for 30 days to induce a wave of ZsGreen-positive RTE, and the mice then maintained on normal chow for the next 21 days to allow RTE egress, homing to, and maturation in, SLO (**Figure 1A**). Therefore, compared to other prior studies, RTE in this manuscript are “older” by up to 4 weeks relative to their time of generation in the thymus, which should be considered when comparing data between studies.

**Figure 1.**
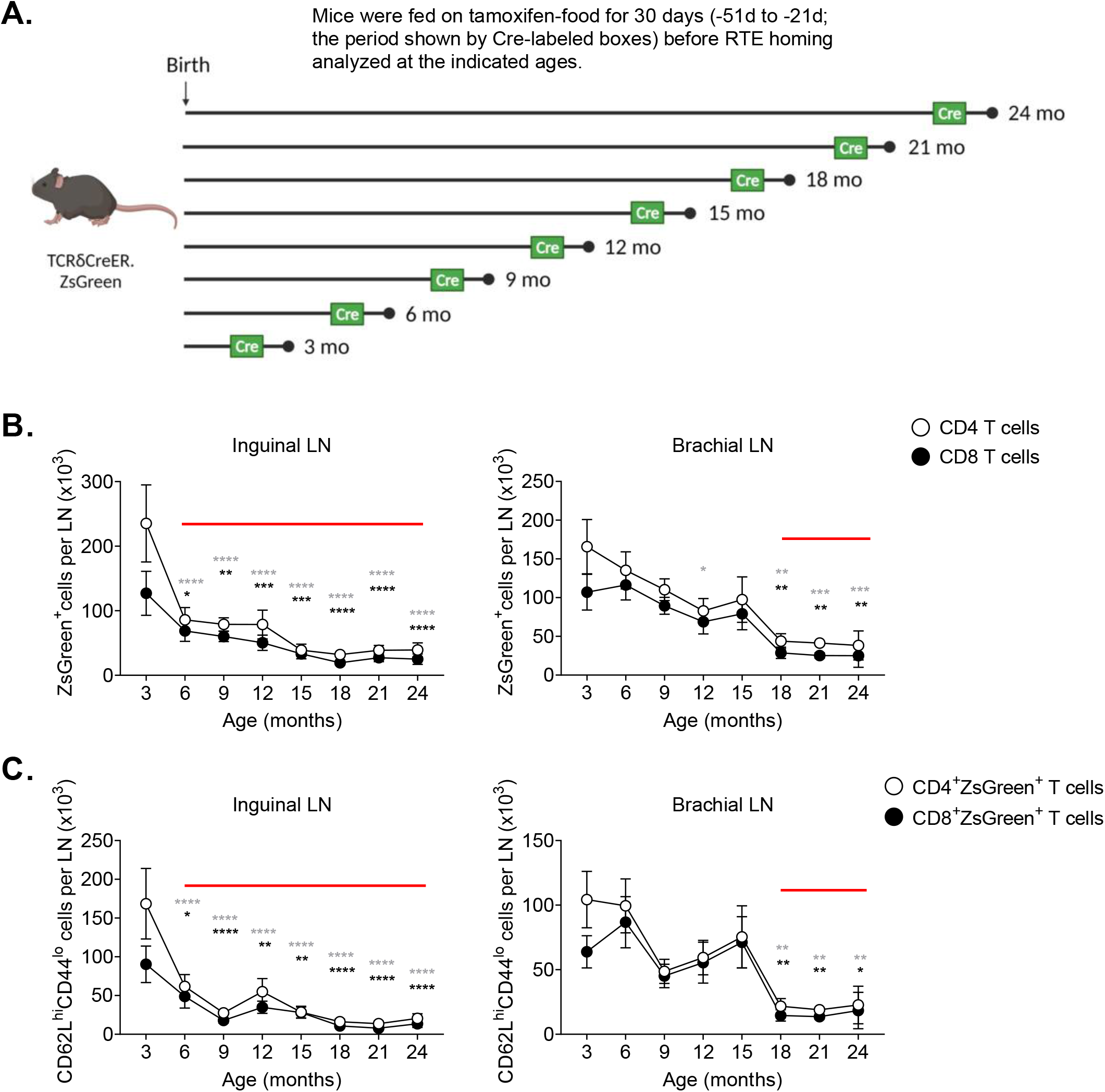
Differential kinetics of seeding and retention of newly generated recent thymic emigrants into the SLO. **(A) (A)** Pictorial representation of experimental strategy to label RTE at different ages, wherein TCRδ^*CreER*^.ZsGreen mice were fed on TAM chow for 30 days and revert back to normal chow for another 21 days before analyzing the ZsGreen^+^ RTE at indicated timeline ages. Phenotypic analysis of cells from SLO were performed using multicolor FCM. Data show absolute numbers of **(B)** ZsGreen^+^ and **(C)** ZsGreen^+^CD62L^hi^CD44^lo^ naive phenotype RTE among the CD4^+^ and CD8^+^ compartments of inguinal and brachial LN. Data represent pooled results of longitudinal experiment performed across 5-6 independent harvests with 9-19 mice/age group **(A-C)**. Error bar represents mean ± SEM **(B, C)**. * p < 0.05, ** p < 0.01, *** p < 0.001, **** p ≤ 0.0001 (p-values for CD4^+^ and CD8^+^ T cells are denoted by grey and black stars, respectively); Two-way ANOVA followed by Dunnett’s multiple comparison test (multiple comparisons to 3 mo). **(B, C)**. The horizontal red line in the graph denotes the time where significant drop in the number of RTE observed.

We first performed a time-course analysis in a cross-sectional manner to evaluate seeding, maturation, and retention of ZsGreen^+^ RTE in different SLO at 3, 6, 9, 12, 15, 18, 21, and 24 mo of age **(Figure S1A)**. In young animals, nearly a third of all cells in SLO were RTE. That proportion dropped to 2-5% in 24-mo old mice. Nonetheless, in agreement with the data from both the Rag2pGFP transgenic mice^20^ and older humans^27^, we detected both CD4 and CD8 RTE even at 24 mo of age **(Fig. S1A, B)**. We found no difference in the ability of RTE across lifespan to mature (lose the surface expression of CD24 and gain Qa-2^4^) once seeded in different SLO (**Fig. S1C**). The ratio of RTE (ZsGreen^+^) to resident, naïve T cells (ZsGreen^-^, CD44^lo^62L^hi^) remained relatively constant across ages, with some variability at 6 and 9 months, and a late decline at 24 months **(Fig. S1D**). The CD4/CD8 ratio of RTE produced at different ages also did not change much over lifespan **(Fig. S1E)**, consistent with the idea that the late-life increase in CD8 T cells over CD4 T cells is a function of differential peripheral maintenance^20^. Moreover, consistent with prior observations from Fink’s group^20^, we found that the production of ZsGreen^+^ single positive cells in the thymus remained constant with age when quantified as a ratio of RTE to CD4^+^CD8^+^ double-positive (DP) thymocytes to control for age-related thymic involution **(Figure S1F)**.

However, major age-related differences were found in the numbers of RTE detected in different SLO. Specifically, skin-draining superficial axillary and inguinal LNs exhibited signs of early numerical decline in RTE around 6-9 mo of age, which further progressed with aging **(Fig. 1B and S2A)**. By contrast, an early decline was not evident in the spleen or in brachial LN, which reside adjacent to the scapula, attached to the biceps muscle, and drain deeper regions of the forearm, shoulder and neck and not the skin^28, 29^. Both the spleen and the brachial LN only showed significant decline in RTE around 18 mo **(Fig. 1B and S2A**); a similar late reduction in RTE was detected in the blood **(Fig. S2A)**.

The age-dependent decline of RTE in the SLO was largely due to the reduction of T_N_ ZsGreen^+^ cells (CD62L^hi^CD44^lo^, **Figure 1C and S2B)**, whereas the numbers of central memory (T_CM_, CD62L^hi^CD44^hi^) and effector memory (T_EM_, CD62L^lo^CD44^hi^) ZsGreen^+^ T cells were very low in all LN, and higher in the spleen **(Figure S1G, H)**. While in the LN both the CD4 and CD8 T_N_ ZsGreen^+^ cells showed the same kinetics of decline (early in the superficial and delayed in deeper LN), we found that splenic (and to a lesser extent, blood) CD4, but not CD8, T_N_ cells showed evidence of an earlier numerical decline **(Figure 1C and S2B)**. RTE of T_N_ phenotype in circulation only showed a significant drop at 21 mo and later **(Figure S2B)**.

These results collectively suggest that while the immediate (1h) immigration into SLO and transition of RTE to mature T_N_ cells seems unaffected by aging, axillary and inguinal LNs experienced an early age-related decline in the numbers of newly generated T cells, starting around 6 mo, whereas for brachial LN and spleen (and, in a limited data set, iliac LN) this defect occurred at the older ages. While it could be argued that this change may be precipitated by the decline in thymic production, the uneven nature of SLO atrophy suggests the action of other mechanisms.

### Decline of CCR7 expression and transient increases of S1P1 correlate with the kinetics of decline of RTE in SLO

The ability of T cells to reside in the LNs depends on several signals, most notably the expression of the C-C chemokine receptor type 7 (CCR7) and sphingosine 1-phosphate receptor (S1P1) on the surface of RTE. CCR7 regulates LN homing and retention, and its high-affinity ligands (CCL19 and CCL21) are produced by LN stromal cells, predominantly FRCs, while S1P1 promotes the egress of T cells from the LNs via efferent lymphatics, with LECs producing its ligand, sphingosine 1-phosphate^30^. We examined discrete waves of reporter^+^ RTE in SLO and found that peripheral LNs and spleen had significantly reduced numbers of CCR7-expressing reporter^+^CD44^-^ naive cells both in CD4^+^ and CD8^+^ compartments starting from 6 mo of age **(Figure 2A and S3A, B)**. This decline in CCR7-expressing RTE was an attribute of SLO as we failed to find changes in the circulation **(Figure S3A, B)**. Interestingly, analysis of CCR7-expression on T_CM_ cells revealed a similar pattern of decline of CCR7^+^CD44^+^ZsGreen^+^CD4^+^ and CCR7^+^CD44^+^ ZsGreen ^+^CD8^+^ cells, with earlier significant decline in the axillary and inguinal LNs as compared to brachial ones **(Figure 2B and S3A, C)**.

**Figure 2.**
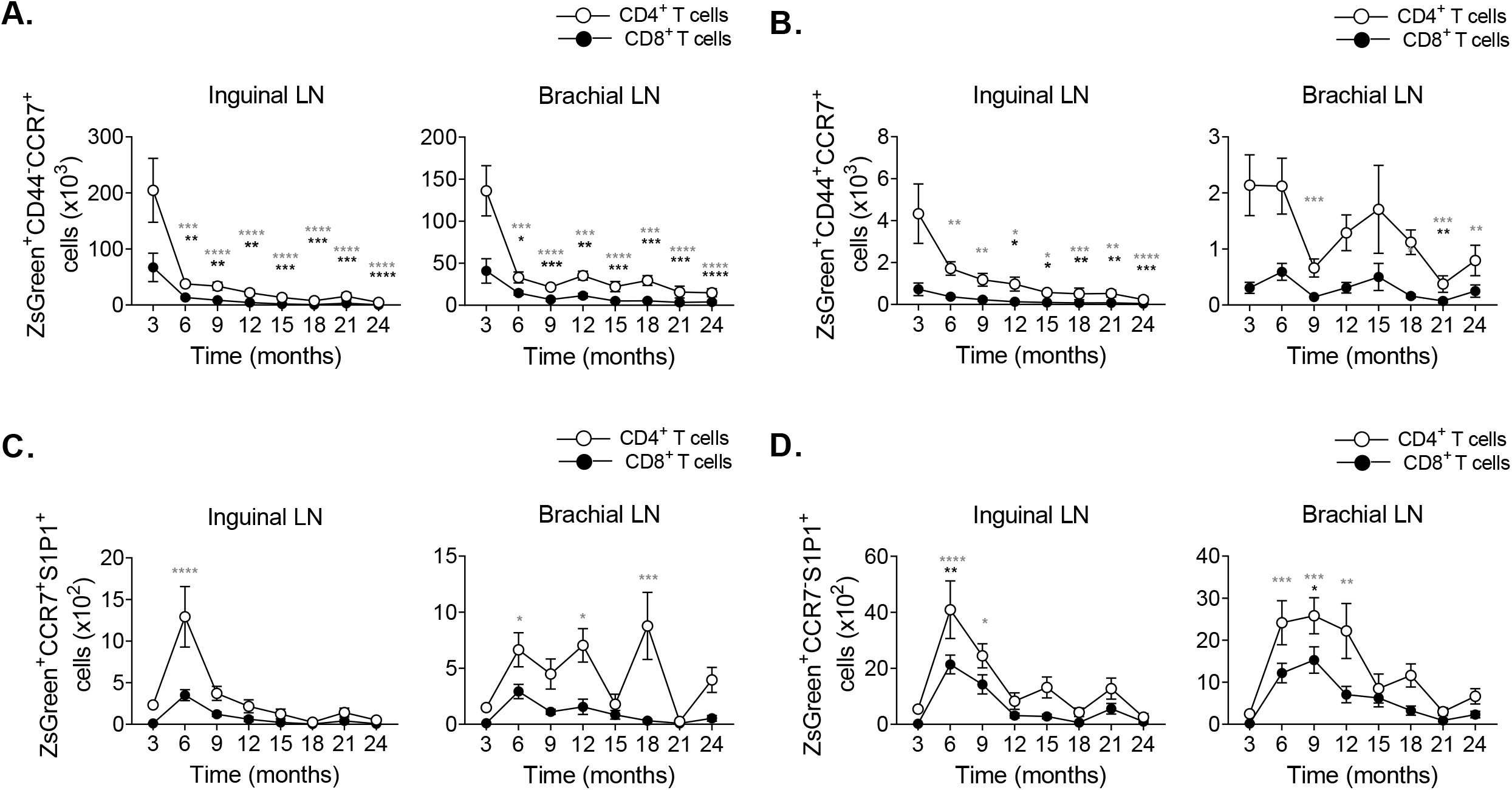
Decline of CCR7-expressing cells and transient increases in S1P1+ RTE correlates with age-related decline of RTE in SLO. TCRδ^*CreER*^.ZsGreen mice were treated as in figure 1A. Peripheral LN, spleen and blood cells were stained with CCR7 and S1P1 and analyzed by FCM. **(A, B)** Data show absolute numbers of **(A)** ZsGreen^+^CD44^-^CCR7^+^ cells and **(B)** ZsGreen^+^CD44^+^CCR7^+^ cells from CD4 (open circle) and CD8 (closed circle) T cell compartment in the inguinal and brachial LN. **(C, D)** Absolute numbers of **(C)** ZsGreen^+^CCR7^+^S1P1^+^ cells and **(D)** ZsGreen^+^CCR7^-^S1P1^+^ cells from CD4 (open circle) and CD8 (closed circle) T cell compartment of the SLO and circulation were shown. Data represent pooled results of longitudinal experiment performed across 5-6 independent harvests with 9-19 mice/age group **(A-D)**. Error bars represent mean±SEM. * p < 0.05, ** p < 0.01, *** p < 0.001, **** p ≤ 0.0001 (p-values for CD4^+^ and CD8^+^ T cells are denoted by grey and black stars, respectively); Two-way ANOVA followed by Dunnett’s multiple comparison test (multiple comparisons to 3 mo) **(A-D)**.

We found that the absolute number of ZsGreen^+^CCR7^+^S1P1^+^ and ZsGreen^+^CCR7^-^S1P1^+^ exhibited a spike at 6-9 mo in the SLO, which was more prominent for CD4 T cells **(Figure 2C, 2D and S3D-F)**. Additionally, expression of CCR7 significantly decreased on the surface of CD4^+^ZsGreen^+^ and CD8^+^ZsGreen^+^ cells as early as 6 mo and remained reduced across SLO thereafter both when evaluated as the number of positive cells and as the mean fluorescent intensity of expression **(Figure 2A, B and S4C)**. In contrast, circulating ZsGreen^+^ cells had elevated expression of CCR7 **(Figure S4C)**. We also noted a transiently increased expression of S1P1 on the surface of CD4^+^ZsGreen^+^ and CD8^+^ZsGreen^+^ cells at around 9 mo in SLO, and observed expected higher S1P1 levels on their circulating counterparts **(Figure S4A, B and D)**. Collectively, reduced representation of CCR7^+^ and transiently increased representation of S1P1^+^ cells in the LN as well as the increase of circulating ZsGreen^+^CD44^-^CCR7^+^ and ZsGreen^+^CD44^+^CCR7^+^CD4^+^ and CD8^+^ T cells suggested that the aging LN microenvironment might not be able to retain these cells, perhaps even facilitating their egress.

### Normal homing and selectively reduced RTE retention characterize LN aging and depend on the age of LN and not of T cells

The numerical decline of RTE in SLO with aging could be due to age-related intrinsic T cell defect(s), to age-related change(s) in SLO microenvironment; or both. Given the observed decline in the numbers of RTE in the superficial pLN around 6-9 months, we sought to establish whether and which of the SLO at this age are receptive to incoming RTE. To test this, we flow cytometry (FCM)-sorted young adult (3 mo) RTE from tamoxifen-induced TCRδ^CreER^.TdTomato mice and transferred them into naïve adult (3 mo), middle-age (9 mo) and older (19 mo) C57BL/6 mice. Our analysis showed that reporter^+^ RTE cells homed efficiently into SLO of all three ages 1 hour after transfer **(Figure 3A and S5A)**. However, analysis at 30 days post transfer revealed that old (but not young and middle-aged) inguinal and brachial LN failed to efficiently retain RTE **(Figure 3B and S5B)**. These data indicate that adult RTE can home to old SLO but are not properly maintained. Reporter^+^ cells appeared to be well-retained in the spleen of all ages, and we noted that significantly larger numbers of RTE circulated in the blood in older mice **(Figure S5B)**. Further, we analyzed the phenotypes of the reporter^+^ cells retained in the SLO after 30 days of transfer and found that old inguinal and brachial LN and spleens exhibited reduced numbers of reporter^+^ cells with naïve phenotype, while the majority of reporter^+^ cells in the circulation had activated/memory phenotype **(Figure 3C and S5C)**. The above patterns were also found in axillary LN, although they were not significant 30 days post transfer, likely due to two outlier samples **(Fig S5B)**. We conclude that the inability of old LN to retain RTE is a function of LN, and not T_N_, cell aging.

**Figure 3.**
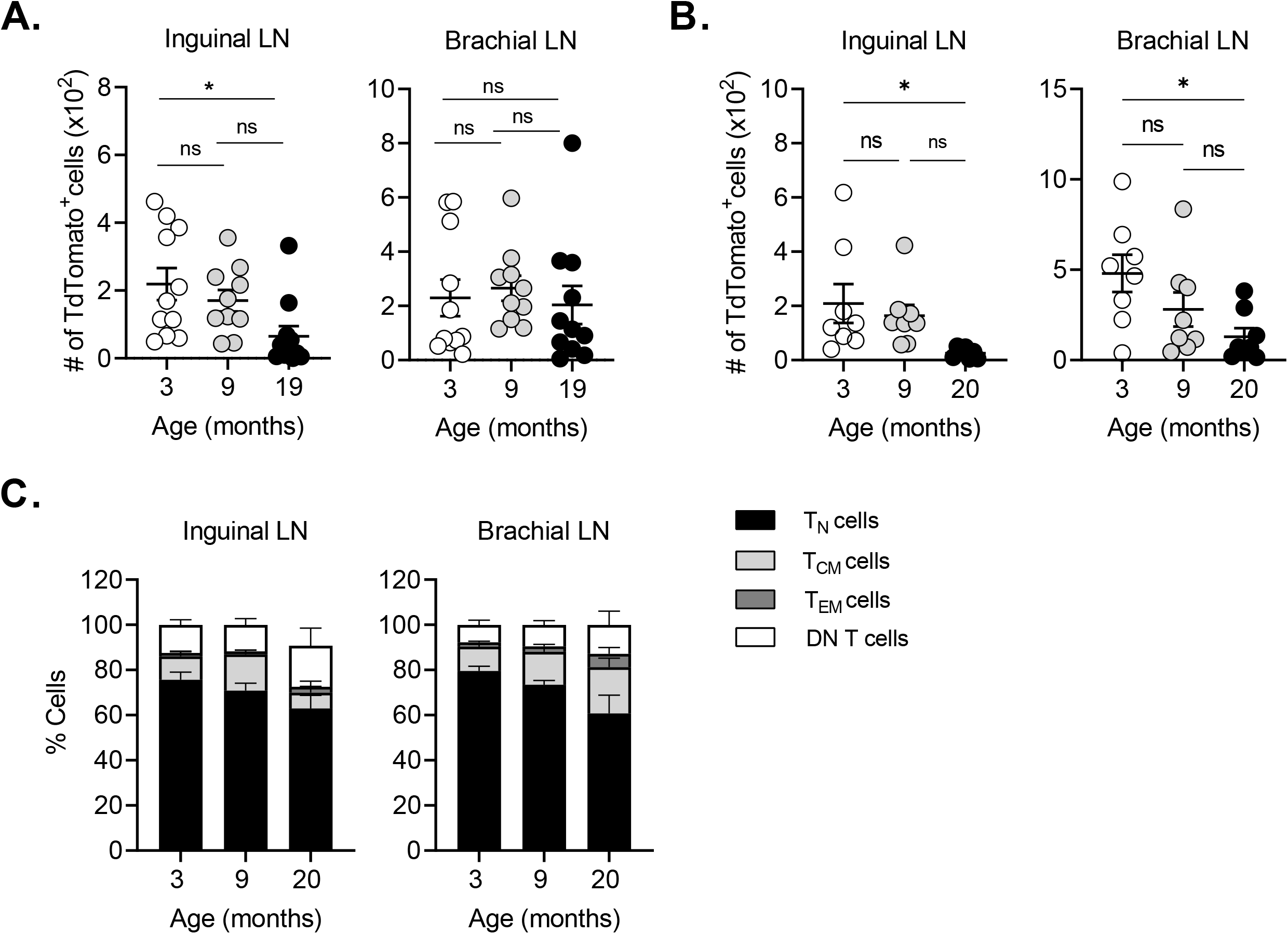
Recent thymic emigrants effectively home to the aged peripheral lymph nodes. TCRδ^CreER^.TdTomato mice were kept on tamoxifen-containing diet for 30 days followed by 21 days on the normal chow before TdTomato^+^ T cells from the spleen and pLN were FACS sorted at 3 mo of age. About 2 × 10^5^ TdTomato^+^ total T cells were *i*.*v*. transferred into 3, 9, and 19 months old naïve C57BL/6 mice. **(A)** One hour later and **(B)** 30 days later, trafficking of transferred cells in the SLO were analyzed. Data show absolute numbers of donor TdTomato^+^ RTE in the inguinal and brachial LN. **(C)** TdTomato^+^ T cells from figure 3B were further analyzed for naïve (CD62L^hi^CD44^lo^), central memory (CD62L^hi^CD44^hi^), effector memory (CD62L^lo^CD44^hi^), and double-negative (DN; CD62L^lo^CD44^lo^) phenotypes and percentages of cell populations among TdTomato^+^ T cells plotted. Data shown are the pool result of two independent transfer experiments with n=8-11 mice per group **(A-C)**. Each dot represents individual mouse **(A-B)**; data represents mean ± SEM **(A-C)**. * p < 0.05, ** p < 0.01, ns = non-significant. One-way ANOVA followed by Tukey’s test **(A, B)**.

### CD4^+^ recent thymic emigrants exhibit increased proliferation and apoptosis in LN with aging

Numerical differences in LN RTE cells could simply reflect their immigration and emigration, or could also involve changes in their proliferation and/or survival. To address this issue, we analyzed RTE proliferation by FCM using BrdU incorporation and Ki-67 expression, and apoptosis using Annexin-V and live/dead-Zombie Aqua staining. Since we observed a significant decline of RTE in pLN around 9 mo, we analyzed time-stamped cells (generated as in **Fig. 1A**) at 3, 9, and 22-23 mo of age. Our analysis revealed increased proliferation of RTE marked by TdTomato^+^ reporter in pLN of 22 mo mice compared to 3 mo adult group, which was particularly pronounced for BrdU^+^Ki67^+^ RTE **(Figure S6A**,**B)**. These results were statistically significant in all LN for CD4 T cells **(Figure S6B)**. Apoptosis was even more dramatically increased in CD4 RTE **(Figure S6C, D)**. Importantly, RTE (reporter^+^) cells in pLN did not show changes in their proliferation or apoptosis at 9 mo compared to 3 mo **(Figure S6A-D)**. Neither the hyper-proliferation nor increased apoptosis of reporter^+^ cells were observed in the spleen at examined time points **(Figure S6B, D)**.

### Age-related changes in the lymph node stromal compartment dominantly impact LEC numbers and the integrity of the FRC reticular networks

As the maintenance of T_N_ cells requires homeostatic and survival signals from the LN stromal cells, we next sought to determine whether the numerical decline of T_N_ cells may correspond to the numerical, structural, and functional changes in LN stromal cells.

Different results have been reported regarding age-related changes in the number LN stromal cells^5, 14, 15^. The disparity may be due to whether the analyses were performed on skin-draining or on deeper, mediastinal LN cells, to the exact enzymatic treatment used for tissue dissociation, to subtle differences in the age of animals, or to other causes. To exclude technical considerations, we performed FCM analysis and enumeration of the pLN (axillary, inguinal, and brachial) stromal cell subsets following digestion with Liberase-TL and DNase-I^15, 19^. Our data show that cells from pooled superficial (axillary and inguinal) LN exhibit a significant reduction of CD45^-^Ter119^-^ non-hematopoietic stromal cells at 12 mo of age and thereafter compared to the 6 mo old mice, although they are not reduced relative to the 3 mo time point **(Figure 4A and 4B)**. The reduction after 6 month was consistent with the pattern of T_N_ cell decline. Furthermore, FRCs (CD45^-^Ter119^-^CD31^-^gp38^+^CD21^-^CD35^-^), LECs (CD45^-^Ter119^-^CD31^+^gp38^+^), and BECs (CD45^-^ Ter119^-^CD31^+^gp38^-^) followed a similar trend of a numerical decline in the axillary and inguinal LN **(Figure 4B)**, with the most pronounced changes in the numbers of LECs. In contrast, the stromal cells in the brachial LN appeared unchanged **(Figure 4B)**. To evaluate another deep tissue-draining LN, we tested iliac LNs that drain the prostate and reproductive organs, and lateral tail. We did not find fluctuations in the stromal cell numbers of the iliac LNs at any of ages analyzed **(Figure S7A)**.

**Figure 4.**
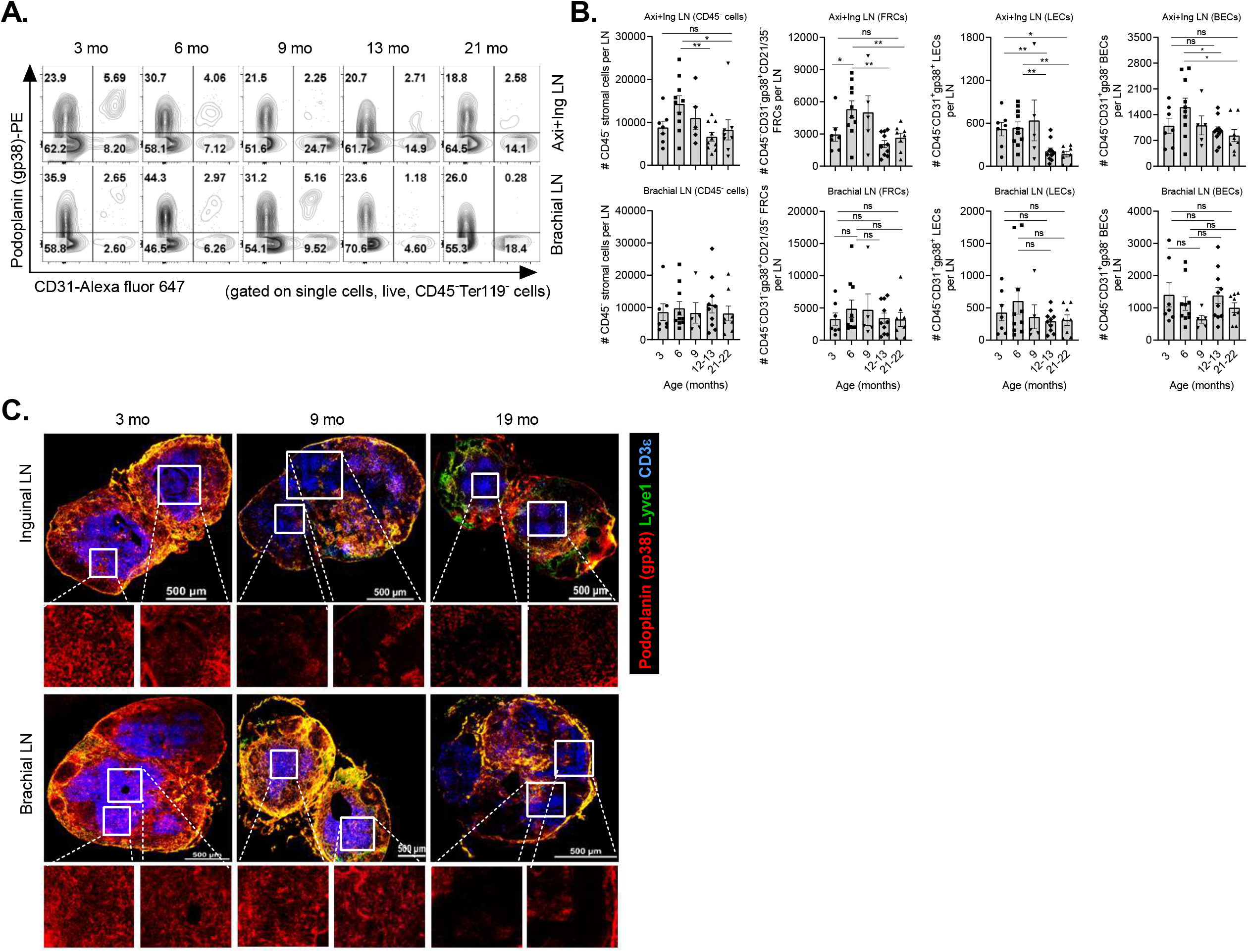
Timing and effect of T_N_ cell numerical decline on the lymph node stromal compartment during aging. One side of peripheral LNs of the naive wild-type C57BL/6 mice of indicated timeline ages were processed for lymphocyte analysis and contralateral ones were digested with Liberase-TL and DNase-I for stromal cell analysis. **(A)** Representative FACS plots show staining of podoplanin (gp38) and CD31 on stromal cells gated on single cells, live, CD45^-^ Terr119^-^ cells. The number in the quadrant indicates frequencies of the indicated cell populations. **(B)** Data show absolute numbers of CD45^-^ total stromal cells, FRCs (CD45^-^CD31^-^gp38^+^CD21^-^ CD35^-^), LECs (CD45^-^CD31^+^gp38^+^), and BECs (CD45^-^CD31^+^gp38^-^) of the pool of axillary and inguinal (Axi+Ing) LN (top) and brachial LN (bottom). Each dot represents individual mouse and data represent mean±SEM. n= 5-10 mice/group. * p < 0.05, ** p < 0.01, *** p < 0.001, **** p ≤ 0.0001, ns = non-significant; One-way ANOVA followed by Dunnett’s multiple comparison test.**(C)** Representative images of 8 μ-thick cryo-sections of the LNs stained with Lyve-1 (green), Podoplanin (gp38; red) and CD3 (blue) were shown. The zoomed images of the region marked with white-square were shown at the bottom of respective images. Images are representative of 5 mice/group. Magnification, 60X; Scale bar, 500 μm.

To interrogate whether the LN stromal networks undergo structural or organizational alterations, we employed confocal microscopy on frozen LN sections at 3, 9, and 19 mo of age. Our image analysis showed that inguinal LN exhibit gross structural alterations in podoplanin (gp38) staining at 9 and 19 mo, suggesting FRC network disruption at these ages **(Figure 4C**, high magnification insets). In parallel, we noted fewer CD3^+^ T cells confined to the characteristic T cell zones at 9 mo, which were further reduced at 19 mo. A similar FRC network disruption was observed at 19 mo in brachial LN, whereas their stromal integrity was well maintained at 9 mo **(Figure 4C)**. We next assessed if FRC network alterations may correlate with the levels of soluble mediators produced by FRC, including the survival factor IL-7 and chemokines CCL19 and CCL21. The IL-7 and CCL21 expression did not change in old LN compared to adult **(Figure S7B)**. Old LN seemed to have more CCL19 compared to adult, however, group means did not reach statistical significance **(Figure S7B)**. Interestingly, CCL19 levels tended to increase wherever the FRC network was perturbed, i.e. at 9 mo in inguinal nodes and 18 mo brachial nodes **(Figure S7B)**. Collectively, this shows that despite unchanged numbers of FRCs at 9 mo, inguinal LN start exhibiting gross FRC network alterations and that similar changes occur in the brachial LN at 19 months, paralleling their reduced receptiveness for RTE **(Fig. 1B, C)**. As this also correlates with the increase in CCL19 in respective LN, it is tempting to speculate that the increase in CCL19 may be compensatory in nature.

### Life-long changes in homeostatic maintenance and phenotype of T cells generated in the early life

How long and how well are T_N_ cells generated in youth maintained in SLO throughout their life? To better understand this, we used TCRδ^CreER^.TdTomato mice, induced by tamoxifen diet for 30 days at 2 months of age, generating a wave of RTE marked for life **(Figure 5A)**. Tracking ‘time-stamped’ cells allowed us to longitudinally monitor and analyze T cells generated at 2 mo, across the life of a mouse. Time-stamped cells generated at 2-3 mo started to decline as early as 5-6 mo of age in the SLO, followed by further, more gradual, decline from 6-24 mo **(Figure 5B and S8A)**. Notably, all SLO showed similar pattern of decline for both CD4^+^ and CD8^+^ time-stamped cells. In the blood, the decline was evident from 12 months of age **(Figure S8A)**. Again, the decline in the overall numbers of cells “born” in youth was driven by a significant decline of T_N_ cells **(Figure 5C and S8B)**. Collectively, these results suggest that aging SLO, and particularly pLN, exhibit an early defect in the maintenance of T cells.

**Figure 5.**
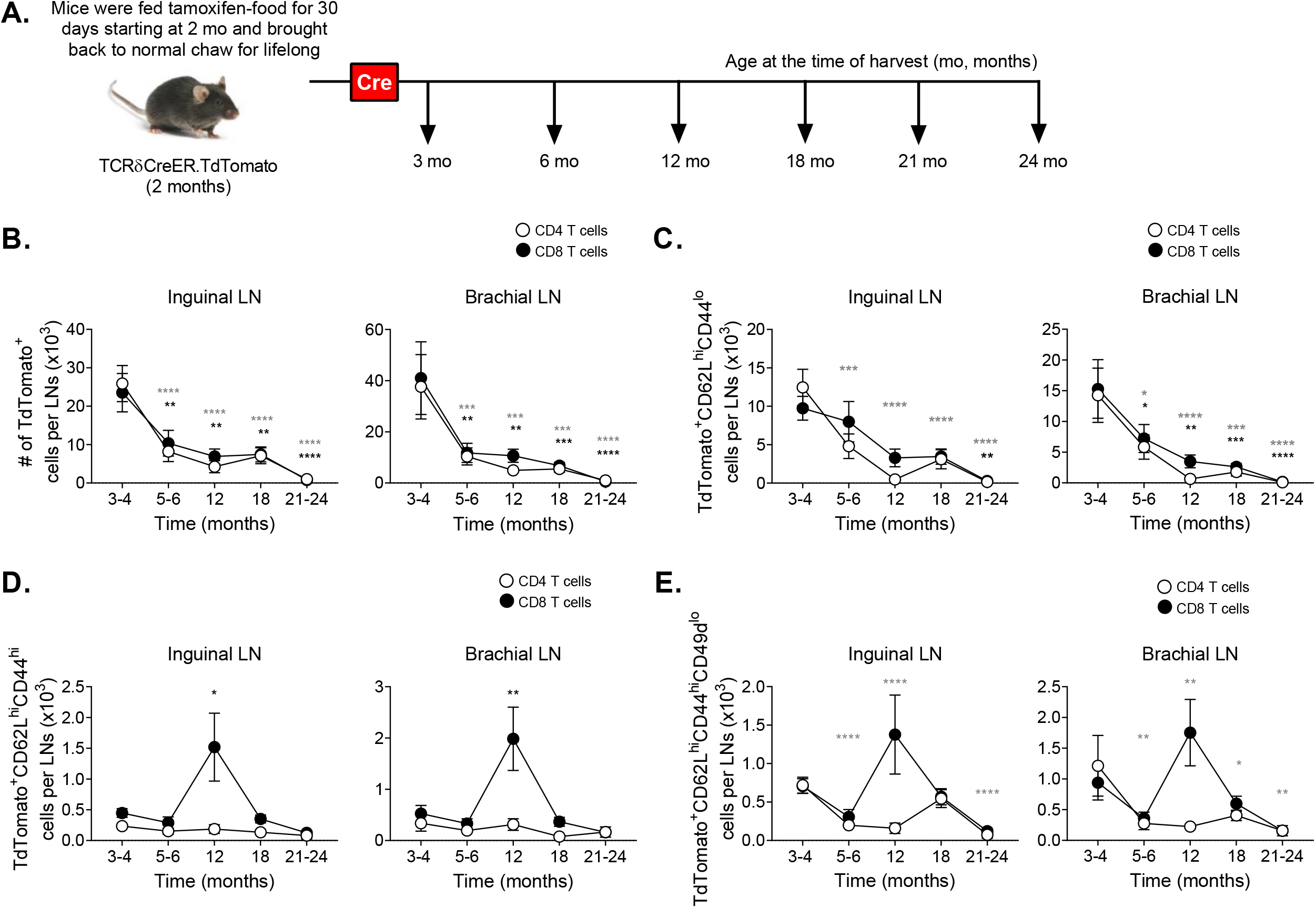
SLO microenvironment failed to maintain T cells generated in the early life during aging under steady-state condition. **(A)** Pictorial representation of experimental strategy to label T cells generated (time-stamped cells) at 2-3 mo of age (time where Cre-expressed), wherein TCRδ^*CreER*^.TdTomato mice were fed on TAM chow for 30 days starting at 2 mo of age and thereafter lifelong maintained on normal chow before analyzing the TdTomato^+^ time-stamped cells by FCM at indicated timeline ages under homeostatic condition. **(B-E)** Absolute numbers of **(B)** total TdTomato^+^ time-stamped cells, and **(C)** TdTomato^+^ time-stamped cells with naive (CD62L^hi^CD44^lo^), **(D)** central memory (CD62L^hi^CD44^hi^) and **(E)** virtual memory (CD62L^hi^CD44^hi^CD49d^lo^) phenotype in the inguinal and brachial LN, spleen, and circulation are shown. Data represent pooled results of longitudinal experiment performed across 5 independent harvests with 11-18 mice/age group **(A-E)**. Error bar represents mean ± SEM. * p < 0.05, ** p < 0.01, *** p < 0.001, **** p ≤ 0.0001 (p-values for CD4^+^ and CD8^+^ T cells are denoted by grey and black stars, respectively); One-way ANOVA followed by Dunnett’s multiple comparison test (compared to 3-4 mo) **(A-E)**.

Some of the CD8^+^ time-stamped cells acquired the T_CM_ (CD62L^hi^CD44^hi^) phenotype, and we detected their spike (a 2-3-fold increase in numbers) at 12 mo and 12-18 mo in SLO and the circulation, respectively, after which they returned to the baseline at older ages **(Figure 5D and S8C)**. While some of the T_N_ cells differentiate into T_EM_ and T_CM_ as a consequence of antigen stimulation, there is also a pool of T_N_ cells that differentiate into T_CM_-like phenotype without exposure to manifest microbial infection^31^, likely due to adaptation to reduced homeostatic signals, and the accumulation of these antigen-independent virtual memory T (T_VM_) cells is one of the hallmarks of aging in specific pathogen-free laboratory mice^32, 33, 34, 35, 36^. We found that most of the accumulated time-stamped CD8^+^ T_CM_ cells were of the virtual memory phenotype (CD62L^hi^CD44^hi^CD49d^lo^) around 12 mo of age in the pLN **(Figure 5E and S8D)**. Moreover, CD8^+^ T_VM_ cells accumulated in the circulation, but not in the spleen **(Figure S8D)**. By contrast, CD62L^hi^CD44^hi^CD49d^hi^ true-central memory T (T_CM_) cells were not detectable in pLN or blood, and did not increase at 12 months in any of the organs examined **(Figure S8E)**.

We conclude that following the initial drop in maintenance of T_N_ cells at an early age, there is a transient wave of conversion of T_N_ cells into T_VM_ cells that is likely compensatory in nature, whereby, presumably, these cells might shift their maintenance pattern from IL-7 to IL-15, consistent with our prior data^33^. In line with this, we observed the increased accumulation of these T_VM_ cells in the bone marrow, a rich source of IL-15, between 6 mo and 18 mo **(Figure S8F)**.

### Participation of RTE in a skin-initiated immune response declines at an early age

Given the surprisingly early changes in superficial LN and in SLO overall, we sought to examine to what extent newly generated T cells (RTE) can participate in an immune response. We challenged tamoxifen-induced TCRδ^CreER^.ZsGreen or TCRδ^CreER^.TdTomato mice with foot-pad injection of RepliVAX-WN (R-WN), a single-cycle WNV vaccine lacking the viral capsid (C) protein, and thus unable to package new virions^37^. R-WN carries the main WNV antigens immunodominant in B6 mice and confers protection against subsequent lethal WNV challenge in both adult and old mice^38^. However, in old mice primary R-WN immunization induces significantly reduced antigen-specific T and B cell responses. We analyzed responses of reporter^+^ cells to WNV-immunodominant NS4b_2488_ CD8 and E_641_ CD4 epitopes using the NS4b_2488_:Db and E_641_:I-A^b^ tetramers^38, 39^ and found robust responses at 3 months of age by percentage and absolute number of tetramer^+^ cells, and differentiation of CD8 cells into cytotoxic granzyme B (GzB)^+^ effectors (**Fig. 6A-D**). However, at 7-8 months, the response had already drastically waned, to the point that the circulating NS4b_2488_:Db tetramer^+^ reporter^+^ cells with a cytotoxic (GzB^+^) phenotype were barely detectable in the blood **(Fig. 6D)**. Virtually identical results have been observed for the CD4 T cell responses against the immunodominant E_641_:I-A^b^ epitope (**Fig. 6E**,**F**). We also examined recall responses in the same animals, and found that, consistent with our prior data^38^, recall stimulation produces strong responses even in the oldest previously vaccinated animals (**Fig. S9A, B, E**) at the level of overall CD8 and CD4 response, particularly in the spleen. Responses by CD4 and CD8 RTE generated at different ages (time-stamped, reporter^+^ cells, **Fig. S9C**,**D**,**F**) were also more numerous and robust, and showed a more gradual pattern of decline, which was particularly evident for the CD8^+^reporter^+^GzB^+^ effector cells in the spleen between 3 and 24 months (**Fig. S9D**).

**Figure 6.**
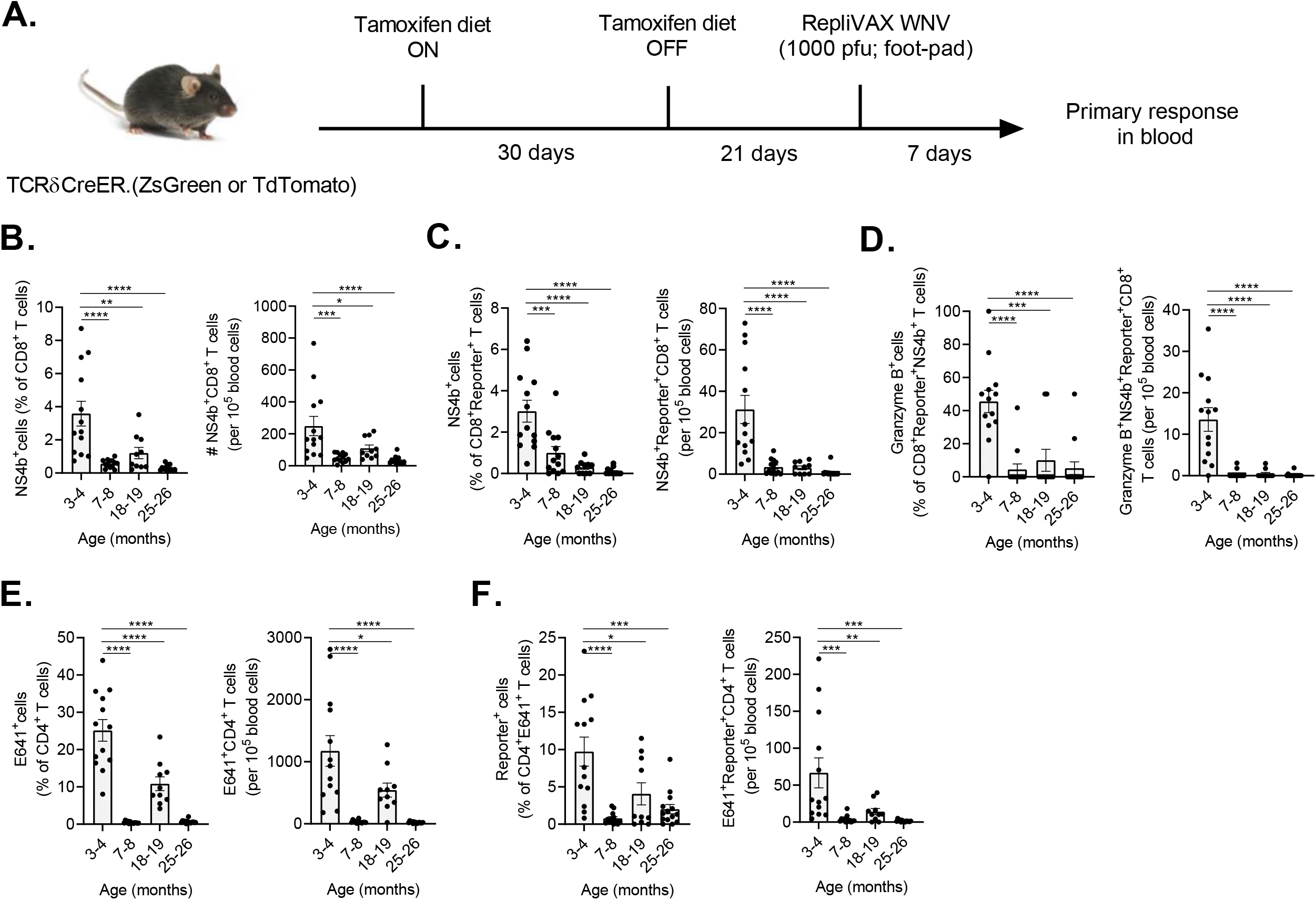
RTE responsive to the Replivax West Nile virus vaccine decline at the age where SLO show defect in RTE maintenance. **(A)** TCRδ^*CreER*^.ZsGreen and TCRδ^*CreER*^.TdTomato mice were fed on tamoxifen-containing diet for 30 days and subsequently maintained on normal diet for next 21 days. At day 21, mice were *s*.*c*. injected at food-pad with Replivax West Nile virus vaccine (1 × 10^3^ pfu). After 7 days’ post injection, blood was collected and West Nile virus specific NS4b^+^ **(B-D)** and E641^+^ **(E, F)** T cells in the reporter (ZsGreen or TdTomato)^+^CD8^+^ and reporter^+^CD4^+^T cell compartment, respectively were analyzed. **(B)** Data show percentages of NS4b^+^ T cells among the CD8^+^ T cells (left) and numbers of circulating NS4b^+^CD8^+^ T cells (right). **(C)** Data show percentages of NS4b^+^ cells among the CD8^+^reporter^+^ T cells (left) and numbers of circulating NS4b^+^reporter^+^CD8^+^ T cells (right). **(D)** Data show percentage of granzyme B^+^ cells among CD8^+^reporter^+^NS4b^+^ T cells (left) and numbers of circulating granzyme B^+^NS4b^+^reporter^+^CD8^+^ T cells (right). **(E)** Percentages of E641^+^ T cells among the CD4^+^ T cells (left) and numbers of circulating E641^+^CD4^+^ T cells (right) were shown. **(F)** Percentages of E641^+^ cells among the CD4^+^reporter^+^ T cells (left) and numbers of circulating E641^+^reporter^+^CD4^+^ T cells (right) were shown. Data represent pooled results from three separate experiments with 10-14 mice per age group **(B-F)**. Each circle represents individual mice. Error bar represents mean ± SEM. * p < 0.05, ** p < 0.01, *** p < 0.001, **** p ≤ 0.0001; One-way ANOVA followed by Tukey’s multiple comparison test **(B-F)**.

Collectively, these results suggest that RTE participate in primary immune responses to subcutaneous/intradermal vaccination at young age but that primary responses are barely detectable as early as 6-8 mo, although they could be expanded by recall stimulation, in which case they wane only later in life. The implications of these findings for immunity and vaccination are discussed below.

## DISCUSSION

Homeostasis of T_N_ cells is maintained by a critical balance between new T cell (RTE) production and entry into SLO and the turn-over (loss) of mature T_N_ cells due to antigenic stimulation (phenotypic conversion into memory cells) or impaired maintenance in the SLO (death or conversion into virtual memory cells)^20^. The numerical loss of T_N_ cell numbers is the most reliable measure of immune aging and is due to the combined effects of an early-life thymic involution that curtails new T_N_ production and a late-life dysregulation of their homeostatic maintenance in secondary lymphoid organs^40^. The main unexpected finding of this study is that superficial skin-draining LN exhibit an early life (6-9 month) defect in T_N_ cell maintenance, while spleen and deeper brachial (and in limited experiments, iliac) LN show late-life maintenance defects (at 18 months and older). This defect corresponded closely to the reduction of LN-homing CCR7^+^ cells and a transient increase in CCR7^-^S1P1^+^ LN emigrants, and to the age-related structural deterioration of the LN FRC network. Further, T cells generated in early life started declining in SLO at 6 mo, and showed signs of phenotypic change towards virtual memory-like phenotype. Finally, functional participation of RTE in cutaneous priming responses also fell precipitously around 6-8 months as measured by effector responses of time-stamped CD4 and CD8 cells against RepliVAX-WNV vaccination.

To integrate these results into a plausible model of lifespan-related events, one needs to consider several points. Previous reports have clearly shown that thymic output in mice start decreasing from 6-7 weeks of life^20, 41^, and our collaborators’ data suggests an even earlier alteration. However, the data presented here cannot be explained by a mere decline in thymic output, because we find that time-stamped RTE in the same mouse at 6-9 months fail to inhabit axillary and inguinal LN, but not brachial LN and spleen, and that the same cells are replete in the blood at the same time points. This suggested that seeding, and/or retention of RTE in superficial SLO may exhibit early defects with aging. Transfer experiments confirmed that seeding after 1h did not seem different across different SLO, and we found that maturation of RTE into the Qa2^hi^CD24^lo^ phenotype^42, 43^ was uniform as well. We therefore hypothesized that RTE/T_N_ retention differs in the course of early aging in different SLO.

In support of this hypothesis, we found that the timeline of RTE decline corresponded closely with the reduced numbers of time-stamped RTE cells expressing CCR7, a key chemokine receptor needed to guide lymphocytes to the SLO in response to CCL19 and CCL21 gradient. These chemokines are produced by lymphoid stromal cells, particularly FRCs and LECs^44,45^. Our time-stamped RTE also showed a (transient) increase in S1P1 in SLO at 6 months in the early-involuting SLO, and a more continuous presence of these cells in the brachial LN and spleen, suggesting that these cells exhibit a SLO-emigrating phenotype, whereby the transient increase in the level of S1P1 might be enough to overcome the reduced CCR7-mediated retention signal strength^46^. We further found an increase in CCR7^+^ RTE (of both the S1P1^-^ and S1P1^+^ phenotypes) in the circulation at the respective ages. Finally, analysis of time-stamped cells generated early in life (between 1.5-3 months) showed that an early decline in T_N_ retention across SLO was followed by a transient wave of antigen-independent T_VM_ cells, consistent with the conversion of T_N_>T_VM_ cells with aging, which then moved from the LN to bone marrow.

Prior work reported variable changes in SLO stroma with aging, with some authors reporting reduced stromal cell numbers^5, 47^ and others not^15^, and with all groups reporting some extent of disorganization in FRC networks in old mice. As some of the above differences could have been caused by different enzymatic dissociation protocols, we have used the protocol of Masters et al.^15^ to reevaluate this issue by examining SLO of different ages and different anatomic locations. While the brachial LN exhibited stable numbers of CD45^-^ stroma, FRC, LEC and BEC across aging, the early-involuting superficial LN (axillary and inguinal) maintained the numbers of total CD45^-^ stromal cells and of FRC and LEC until 9 months, and then exhibited a significant decline in all three populations from 12 months on. Structural analysis of SLO stroma organization revealed additional important points, showing that a numerically unperturbed stromal compartment does not equate normal structure of stromal networks. Specifically, while the inguinal LN at 9 month and brachial LN at 19 month each contained comparable stromal cell, FRC and LEC numbers, they each exhibited a striking defect in the FRC (podoplanin/gp38^+^Lyve1^-^ cells in the images) network that was both markedly less dense and more diffuse at the sites that correspond to LN T cell zones. This deterioration of the LN FRC network at different times in different LN could be expected to lead to inefficient access of T_N_ cells to chemotactic and maintenance factors, including IL-7. We believe that in this case access, and not protein levels, may be limiting, because, as previously reported^5, 14^, we found that with aging, levels of IL-7 itself were not reduced compared to young (and neither were those of CCL19 and CCL21), and in fact we found a potential compensatory increase at 9 months for IL-7 and CCL19.

Based on all of the above, we can integrate the data into a hypothetical model based on the putative crosstalk between lymphocytes and stromal cells in SLO **(Figure 7)**. Step 1 would occur as the initial reduction of RTE/T_N_ from the thymus leads to less trafficking of these cells to the SLO, somewhere around 3-5 months of life in murine superficial SLO. This will result in an initial subtle insult/injury to the SLO stromal cells, perhaps in the form of reduced trophic signals to FRC/LEC. Step 2 would ensue when FRC/LEC react to this reduction by reduced metabolic or functional fitness, and FRC potentially begin to retract their processes and thereby begin to reduce the density of the FRC network. This will result in reduced survival/trophic signaling to the surrounding T_N_ cells (step 3), and the entire sequence will progress as a feed-forward negative loop, leading to increased SLO stromal dysregulation and progressive reduction in RTE/T_N_ maintenance. Consequences of this loop are evident as reduced RTE/T_N_ retention, disrupted FRC networks, changes in RTE/T_N_ homing receptor expression towards emigrating phenotype and subsequently towards virtual memory fate and alternative maintenance. At older ages, we found that this can even take a form of increased proliferation, and, in the case of CD4 T cells, also increased apoptosis, previously described in the literature^20, 48^. It remains to be determined whether CD4^+^ T_N_ cells in old LN are experiencing proliferation-induced apoptosis or may be attempting to compensate for increased apoptosis by increasing the rate of proliferation.

**Figure 7.**
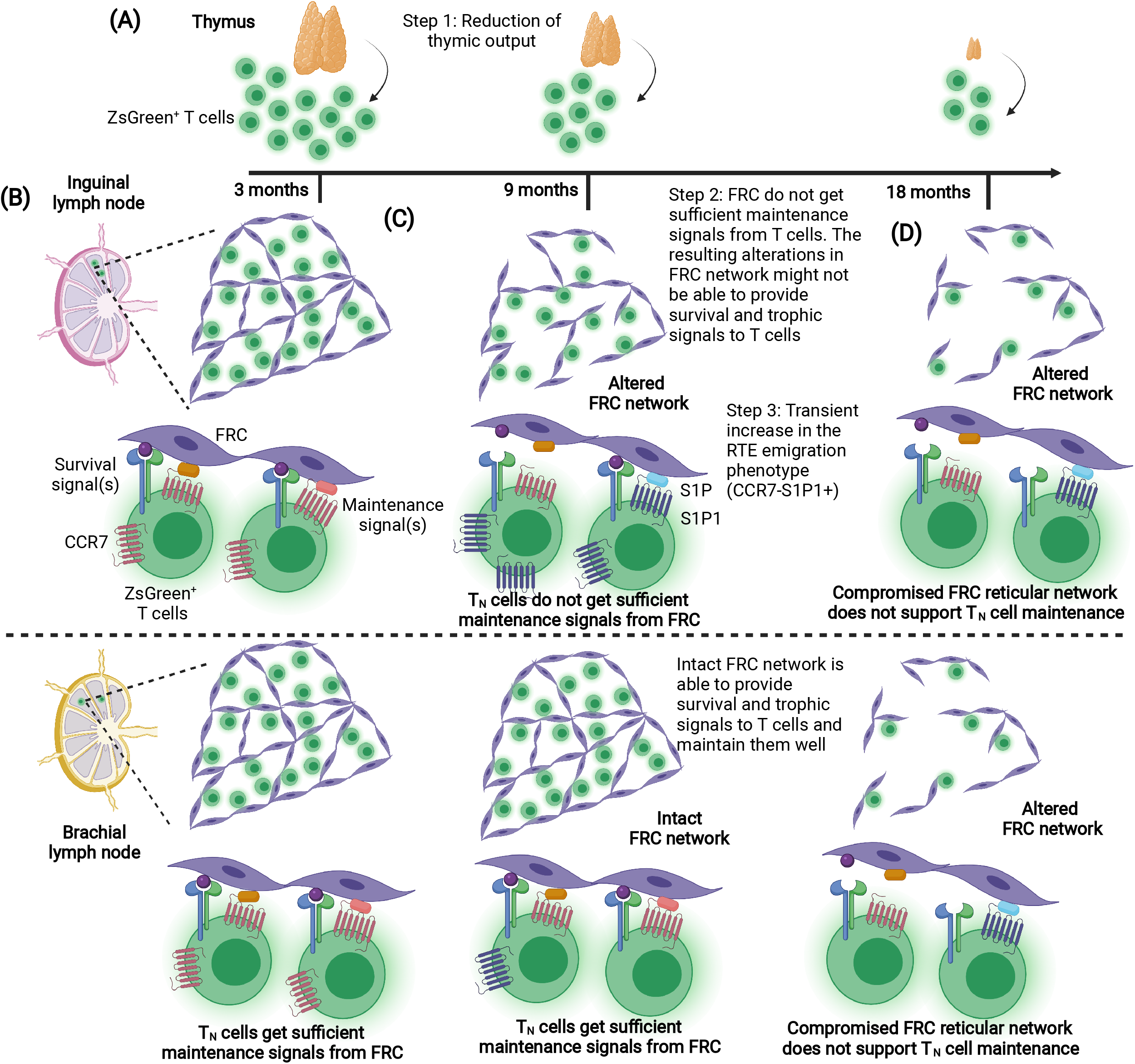
Asymmetric involution of the lymph nodes. Based on the data presented here, we hypothesized a model (Created with BioRender.com), which summarizes the timing of the T_N_ maintenance defect and the extent of SLO damage. **(A)** Reduction of the thymic output (in our case, ZsGreen^+^ T cells) with increasing age leads to the reduction of the seeding of the ZsGreen^+^ T cells (RTE) into the SLO, including inguinal and brachial LN. **(B)** At adult age, around 3 mo, pLN (both inguinal and brachial LN) show intact FRC (gp38/podoplanin) network. The FRC provide optimal survival and tropic signals (such as IL-7, CCL19, and CCL21) to the T cells. Under homeostatic conditions, these T cells maintained well with optimal bi-directional signals being exchanged for the survival and fitness of T cells and FRC. **(C)** However, age-related changes in the inguinal LN but not the brachial LN induce the perturbations in the FRC architecture and organization, which may invoke alterations in the FRC-T cell bi-directional signal exchanges, leading to metabolic and functional alterations in the FRC. These events may provoke T cells to gain alterative maintenance with altered phenotype (such as generation of TVM phenotype or LN emigrating phenotype (CCR7^lo^S1P1^+^), leading to age-related increased accumulation of CCR7-expressing T cells in the circulation. **(D)** With the advancing age, these events may continue repeatedly, and might be contribute to the deterioration of FRC-T cell bi-directional cross-talk. At around 18-19 mo, the alterations in the FRC network and organization worsen in both the inguinal and brachial LN, and it may no longer supportive for the T cell maintenance, where significant drop in ZsGreen^+^ T cells were seen.

The above model remains to be experimentally tested in greater detail, particularly on the mechanistic side. Work to untangle what mechanisms make superficial, but not deeper, LN susceptible to early atrophy is currently in progress. Regardless of the mechanisms, however, we have shown that the end result of all of the above changes is a drastic compromise of cutaneous immunity to the point that already at 6-8 months of age (an equivalent to early middle-age in humans), superficial LN and the cells therein practically fail to mount any meaningful effector T cell response in laboratory specific pathogen-free mice. Practical considerations here will need to be further investigated as related to older humans, but these findings certainly correspond very well to clinical observations that older adults rarely, if ever, react to acute infection with subcutaneous lymphadenopathy^17, 49, 50^.

## MATERIALS AND METHODS

### Mice

Wild-type C57BL/6 mice of varying ages were obtained from the National Institute of Aging breeding colony. B6.129S-Tcrd^*tm1*.*1(cre/ERT2)Zhu*^/J (TCRδ^*CreER*^), B6.Cg-*Gt(ROSA)26Sor*^*tm6 (CAG-ZsGreen1)Hze*^/J (Rosa26.ZsGreen), and B6.Cg-*Gt(ROSA)26Sor*^*tm14(CAG-TdTomato)Hze*^/J (Rosa26.TdTomato) mice were purchased from the Jackson Laboratory (Bar Harbor, ME). TCRδ^*CreER*^ mice were bred either with Rosa26.ZsGreen or Rosa26.TdTomato to generate TCRδ^*CreER*^.ZsGreen and TCRδ^*CreER*^.TdTomato reporter strains, respectively. All mice strains were maintained under specific pathogen-free condition at the University Animal Care facility, University of Arizona, Tucson, AZ. All experiments were performed in compliance with the guidelines of the University of Arizona’s Institutional Animal Care and Use Committee (IACUC).

### Mice treatment

TCRδ^*CreER*^.ZsGreen mice were fed Tamoxifen (TAM)-containing chow for 30 days followed by normal chow for the next 21 days before they were euthanized to analyzed RTE at 3, 6, 9, 12, 15, 18, 21, and 24 mo of age. Alternatively, TCRδ^*CreER*^.TdTomato mice at 2 mo of age were fed on TAM chow for 30 days followed by lifelong maintenance on normal chow.

### Lymph node stromal cell isolation and analysis

For simultaneous analysis of stromal cells and lymphocytes, left side of the axillary, inguinal and brachial LN were processed for stromal cell analysis and right side were used for lymphocyte enumeration. LN stromal cells were isolated as described earlier^15, 51^ with some modifications. Briefly, a pool of axillary and inguinal, or individual brachial or iliac LNs were digested in a 5 ml transparent U-bottom FACS tube with 1.0 ml of digestion buffer containing 0.2 mg/ml Liberase-TL (Sigma, Saint Louis, Mo) and 20 μg/ml DNase-I (Sigma) in RPMI-1640. Tubes were incubated at 37°C water bath for 20 minutes with continuous shaking at 250 rpm. Samples were pipetted several times using a 1.0 ml pipette to dissociate the LN capsule, and incubation continued further for 20 minutes, followed by vigorous pipetting using 1.0 ml pipettes to dissociate LN tissue. Tubes were further incubated at 37°C water bath for 20 minutes. This resulted in total 60 minutes digestion, at the end of which complete dissociation of LN tissue was evident from the visible lack of LN tissue pieces. Cells were passed through 100 μm cell strainer, and washed with RPMI-1640 containing 10% FBS (Omega Scientific, CA, USA). Cell viability was determined by Trypan blue dye exclusion test, which consistently found >95% viable cells that were counted and processed with flow cytometry staining. Briefly, about 4 × 10^6^ cells were stained with 1 μl mouse Fc-block (anti-CD16/32; BioLegend, San Diego, CA) in 50 μl FACS buffer (PBS containing 2% FBS) for 20 minutes at 4°C in the dark, followed by staining with a cocktail of antibodies (1:100 dilutions of each of anti-mouse CD45-eFluor 450, anti-mouse Ter119-eFluor 450, anti-mouse CD31-Alexa Fluor 647, and anti-mouse CD21/CD35-APC/Cy7 and 1:50 dilution of anti-mouse podoplanin (gp38)-PE) in 50 μl FACS buffer for 60 minutes at 4°C in the dark. Cells were washed with FACS buffer followed by PBS wash and stained with 100 μl of 1:1000 dilution of fixable Zombie Aqua (BioLegend, San Diego, CA) dye for 20 minutes at 4°C in dark. Cells were washed with PBS, fixed with BD Cytofix™ fixation buffer for 5 minutes, washed twice with FACS buffer, and resuspended in 50 μl FACS buffer containing 20 μl CountBright absolute counting beads (Invitrogen, Waltham, MA). Samples were acquired on LSR FoRTEsa (BD Biosciences) and data were analyzed using FlowJo v10.6.1 (TreeStar, Ashland, OR).

### Flow cytometry

Cells were harvested from the peripheral lymph nodes, thymus, and spleen via pressing them through a 40 μm cell strainer using a syringe plunger. Spleen cells and blood were treated with ACK lysis buffer to lyse RBCs. About 2-4 × 10^6^ cells were stained with 1 μl mouse Fc-block (anti-CD16/32) in 50 μl FACS buffer (PBS containing 2% FBS) for 20 minutes at 4°C in dark, followed by staining with surface antibodies (for detailed list of antibodies are given in Table S1) in 50 μl FACS buffer for 40 minutes at 4°C in dark. Alternately, to analyze the expression of homing receptors, CCR7 and S1P1 on RTE, we stained cells with 1 μg each of anti-mouse CCR7-PE/Cy7 and anti-mouse S1P1 (EDG-1)-APC at 37°C for 40 minutes in dark. Cells were washed and stained with 100 μl of 1:1000 dilution of fixable Zombie Aqua (BioLegend, San Diego, CA) for 20 minutes at 4°C in dark, fixed with BD Cytofix™ fixation buffer for 5 minutes, washed twice with FACS buffer, and resuspended in FACS buffer containing 20 μl CountBright absolute counting beads (Invitrogen). Fluorescence minus one (FMO) controls were performed routinely and are shown in some supplemental figures (e.g. in **Figs. S3A** and **S3D** for CCR7 and S1P1). Samples were acquired on LSR Fortessa (BD Biosciences) and data were analyzed using FlowJo v10.6.1 (TreeStar, Ashland, OR). Absolute number of cells were determined as; Cells/μl = [((cell count / counting bead count) x (counting bead volume / cell volume)) x counting bead concentration (beads/μl)].

### Immunohistochemical staining of lymph nodes

Mice were euthanized via isoflurane overdose, peripheral LN were harvested, and immediately snap-frozen in OCT medium (Sakura Finetek, Torrance, CA). 8-10 μm cryo-sections were prepared and fixed in chilled acetone for 5 minutes, air dried, washed with PBS, and treated with 0.3% hydrogen peroxide for 60 minutes at room temperature (RT), washed with ice-cold PBS, followed by blocking with 10% goat serum (Invitrogen) for 60 minutes at RT. Sections were incubated with primary antibodies (1:200 dilutions each of rabbit anti-mouse Lyve-1, syrian hamster anti-mouse podoplanin (gp38)-biotin, rat anti-mouse CD3-APC) for overnight (12-14 hours) at 4°C in the dark, followed by washing four times with ice-cold PBS and incubated with fluorochrome-conjugated secondary antibody or reagents (1:1000 dilution) for 60 minutes at RT in the dark. Sections were washed four times with PBS, fixed with 1% paraformaldehyde (PFA) for 5 minutes at RT, and mounted with Prolong Gold anti-fade mounting reagent (Invitrogen). Tyramide signal amplification kit was used to reveal the podoplanin (gp38) staining according to manufacturer’s instructions. Sections were visualized under Leica DMI6000 inverted fluorescent microscope (Leica Microsystems, Germany) or Nikon Eclipse Ti2-E inverted microscope with a Crest X-Light V2 L-FOV spinning disk confocal and a photometrics prime 95B-25MM back-illuminated sCMOS camera using plan apochromat 60X (Oil/1.4) objective (Nikon Instruments Inc., Melville, NY). Images were analyzed using LAS-X software (Leica) or NIS-Elements imaging software (Nikon Instruments Inc.) at University of Arizona Cancer Center’s imaging core.

### Apoptosis of recent thymic emigrants

TCRδ^*CreER*^.ZsGreen and TCRδ^*CreER*^.TdTomato mice were fed with TAM chow for 30 days followed by normal chow for 21 days before ZsGreen^+^ or TdTomato^+^ RTE were analyzed at 3, 9, and 21 mo of age. 2 × 10^5^ cells from the lymph node and spleen were surface stained with anti-mouse CD4-Brilliant Violet 750 and anti-mouse CD8-Brilliant Violet 785 for 30 minutes at 4°C in dark, stained with Zombie Aqua fixable viability kit (BioLegend), washed and stained with Annexin-V-APC as described previously^52^. Apoptosis and death of RTE were determined by analyzing frequencies of Annexin-V^+^Zombie Aqua^-^ and Annexin-V^-^Zombie Aqua^+^ cells amongst CD8^+^ZsGreen^+^ or CD8^+^TdTomato^+^ cells.

### Proliferation of recent thymic emigrants

TCRδ^*CreER*^.ZsGreen or TCRδ^*CreER*^.TdTomato mice were fed with TAM chow for 30 days followed by normal chow and 1 mg/ml 5-bromo-2’-deoxyuridine (BrdU, Sigma, St. Louis, MO) in drinking water containing 1% glucose (to reduce the taste aversion and increase the palatability of water) for 21 days. BrdU water prepared in sterile water, protected from light and changed every other day. The ZsGreen^+^ or TdTomato^+^ RTE were analyzed at 3, 9, and 21 mo of age. BrdU staining was performed as described previously^52^. Briefly, a total of 5 × 10^6^ lymph node and spleen cells were surface stained with anti-mouse CD4-Brilliant Violet 750 and anti-mouse CD8-Brilliant Violet 785 for 30 minutes at 4°C in dark, washed twice with FACS buffer, stained with Zombie Aqua fixable viability kit (BioLegend), followed by fixation and permeabilization with BD Cytofix/Cytoperm™ solution kit. Permeabilized cells were incubated with 500 Kunits U/ml DNase-I, and stained with anti-BrdU-Alexa Fluor 647 and anti-mouse Ki-67-Brilliant Violet 650 at 4°C for 45 minutes in dark. Cells were analyzed on LSR Fortessa (BD Biosciences) and data were analyzed using FlowJo v10.6.1 (TreeStar, Ashland, OR).

### In vivo homing of recent thymic emigrants (RTE) to secondary lymphoid organs

TCRδ^*CreER*^.ZsGreen and TCRδ^*CreER*^.TdTomato mice were fed with TAM chow for 30 days followed by normal chow for 21 days before mice were euthanized at 3 mo of age. Spleen and pLN were harvested, single cell suspensions prepared, and total T cells were enriched using mouse CD3ε Microbead kit (Miltenyi Biotec), according to manufacturer’s instructions. RTE (ZsGreen^+^ or TdTomato^+^) were FACS sorted, and a total of 2 × 10^5^ cells were *i*.*v*. transferred into 3 mo, 9 mo, and 19-20 mo old C56BL/6 mice via retro-orbital route. The homing or retention of reporter-positive RTE in the SLO were analyzed by flow cytometry after 1 hour and 30 days post transfer.

### IL-7, CCL19 and CCL21 ELISA

A pool of axillary and inguinal LN, brachial LN were homogenized in 350 μl of tissue lysis buffer comprising of 0.5% NP-40 in PBS and protease inhibitor cocktail (Sigma), allowed to lyse at RT for 30 min, centrifuged at 1000g for 15 minutes. Supernatants were immediately frozen at −80°C until analysis. The IL-7, CCL19, and CCL21 were quantified as described previously^15^ using mouse IL-7, mouse CCL21, and mouse CCL19 DuoSet ELISA (R&D Systems) as per manufacturer’s instructions. The IL-7, CCL19, and CCL21 levels were normalized to total protein concentration of the homogenates determined using Pierce BCA protein Assay kit (ThermoFisher).

### Analysis of RTE responses to West Nile vaccination

TCRδ^*CreER*^.TdTomato or TCRδ^*CreER*^.ZsGreen mice were fed on tamoxifen-containing diet for 30 days and subsequently maintained on normal diet for next 21 days. At day 21, mice were *s*.*c*. injected in hind food-pad with Replivax West Nile virus vaccine (1 × 10^3^ pfu). At day 7 post injection, mice were bled retro-orbitally, and primary responses to vaccination were assessed using FCM analysis of WNV-immunodominant NS4b_2488_ CD8 and E_641_ CD4 epitopes using the NS4b_2488_:Db and E_641_:I-A^b^ tetramers. Mice were further received a booster dose of Replivax West Nile virus vaccine (1 × 10^3^ pfu) at day 45 after primary immunization, and recall responses of total and reporter-positive NS4b^+^CD8^+^ T cells and E641^+^CD4^+^ T cell were analyzed in the SLO.

### Statistical analysis

Data for RTE numbers were collected over multiple timeline experiments with overlapping age-groups, and pooled together. Data are expressed as the mean ± SEM. As indicated in the figure legends, statistical analysis was performed by One-way or two-way analysis of variance (ANOVA) followed by Dunnet’s multiple comparison test and Mann-Whitney test using Prism 8 software. Comparisons of the group mean differences were considered significant at p < 0.05.

## Abbreviations

FRC: fibroblastic reticular cells
LEC: lymphatic endothelial cells
pLN: peripheral lymph nodes
RTE: recent thymic emigrants
SLO: secondary lymphoid organs
Tn: naïve T cells

## ACKNOWLEDGEMENT

We would like to thank Dr. Y. Zhuang, Duke University, for generously providing us TCRδCre.ER transgenic mice. We thank Mr. Jose L Padilla-Torres, and University animal care facility, University of Arizona, Tucson, AZ for maintaining mouse colonies. We are grateful to the University of Arizona / UA Cancer Center Shared Flow Cytometry Resource and the Imaging Core Facility, supported by the UACC Core Cancer Center Support Grant P30 CA023074 for technical help with flow cytometry and microscopy. We would like to thank Dr. Laura Hale, The Duke Human Vaccine Institute, Duke University, Durham, NC; Dr. Nancy R. Manley, Department of Genetics, University of Georgia, Athens, GA; Dr. Ellen R. Richie, MD Anderson Cancer Center, Austin, TX; and Dr. Lauren I. R. Ehrlich, Department of Molecular Biosciences, Institute of Cellular and Molecular Biology, The University of Texas at Austin, Austin, TX for scientific discussion, critical reading of manuscript, and inputs.

## FUNDING STATEMENT

This work is supported in part by the USPHS awards AG052359 and AG020719 and the Bowman Endowed Professorship in Medical Sciences (J.N-Ž). Work performed at Duke University was in the Duke Regional Biocontainment Laboratory which received partial support for construction from NIH/NIAID (AI058607, G.D.S).

## CONFLICT OF INTEREST

J.N.Ž. is co-chair of the scientific advisory board of and receives research funding from Young Blood Institute, Inc. YBI, Inc. had no influence on any aspect of this work or the present manuscript.

## AUTHORS CONTRIBUTION

S.A.S., J.L.U., G.D.S., J.A.D., M.R.M. v.d. B., J.N-Ž designed research; S.A.S., J.L.U., C.P.C., M.J. performed research; S.A.S., C.P.C., J. N-Ž analyzed data; S.A.S., J. N-Ž. wrote paper; all authors edited paper; J. N-Ž. Conceived project and supervised the study; G.D.S., J.A.D., M.R.M. v.d. B., J. N-Ž. acquired funding.

## DATA AVAILABILITY

All data available upon request.

## Notes

### Competing Interest Statement

J.N.Z. is co-chair of the scientific advisory board of and receives research funding from Young Blood Institute, Inc. YBI, Inc. had no influence on any aspect of this work or the present manuscript.

## REFERENCES

1. Berzins, S.P., Boyd, R.L. & Miller, J.F. The role of the thymus and recent thymic migrants in the maintenance of the adult peripheral lymphocyte pool. J Exp Med 187, 1839–1848 (1998).

2. Fink, P.J. The biology of recent thymic emigrants. Annu Rev Immunol 31, 31–50 (2013).

3. Berkley, A.M. & Fink, P.J. Cutting edge: CD8+ recent thymic emigrants exhibit increased responses to low-affinity ligands and improved access to peripheral sites of inflammation. J Immunol 193, 3262–3266 (2014).

4. Boursalian, T.E., Golob, J., Soper, D.M., Cooper, C.J. & Fink, P.J. Continued maturation of thymic emigrants in the periphery. Nat Immunol 5, 418–425 (2004).

5. Becklund, B.R. et al. The aged lymphoid tissue environment fails to support naive T cell homeostasis. Sci Rep 6, 30842 (2016).

6. Link, A. et al. Fibroblastic reticular cells in lymph nodes regulate the homeostasis of naive T cells. Nat Immunol 8, 1255–1265 (2007).

7. Knoblich, K. et al. The human lymph node microenvironment unilaterally regulates T-cell activation and differentiation. PLoS Biol 16, e2005046 (2018).

8. Alexandre, Y.O. & Mueller, S.N. Stromal cell networks coordinate immune response generation and maintenance. Immunol Rev 283, 77–85 (2018).

9. Qi, Q., Zhang, D.W., Weyand, C.M. & Goronzy, J.J. Mechanisms shaping the naive T cell repertoire in the elderly - thymic involution or peripheral homeostatic proliferation? Exp Gerontol 54, 71–74 (2014).

10. Yanes, R.E., Gustafson, C.E., Weyand, C.M. & Goronzy, J.J. Lymphocyte generation and population homeostasis throughout life. Semin Hematol 54, 33–38 (2017).

11. Nikolich-Zugich, J. The twilight of immunity: emerging concepts in aging of the immune system. Nat Immunol 19, 10–19 (2018).

12. Goronzy, J.J. & Weyand, C.M. Mechanisms underlying T cell ageing. Nat Rev Immunol 19, 573–583 (2019).

13. Krishnamurty, A.T. & Turley, S.J. Lymph node stromal cells: cartographers of the immune system. Nat Immunol 21, 369–380 (2020).

14. Thompson, H.L. et al. Lymph nodes as barriers to T-cell rejuvenation in aging mice and nonhuman primates. Aging Cell 18, e12865 (2019).

15. Masters, A.R. et al. Assessment of Lymph Node Stromal Cells as an Underlying Factor in Age-Related Immune Impairment. J Gerontol A Biol Sci Med Sci 74, 1734–1743 (2019).

16. Richner, J.M. et al. Age-Dependent Cell Trafficking Defects in Draining Lymph Nodes Impair Adaptive Immunity and Control of West Nile Virus Infection. PLoS Pathog 11, e1005027 (2015).

17. Lazuardi, L. et al. Age-related loss of naive T cells and dysregulation of T-cell/B-cell interactions in human lymph nodes. Immunology 114, 37–43 (2005).

18. Uhrlaub, J.L. et al. Dysregulated TGF-beta Production Underlies the Age-Related Vulnerability to Chikungunya Virus. PLoS Pathog 12, e1005891 (2016).

19. Masters, A.R., Jellison, E.R., Puddington, L., Khanna, K.M. & Haynes, L. Attrition of T Cell Zone Fibroblastic Reticular Cell Number and Function in Aged Spleens. Immunohorizons 2, 155–163 (2018).

20. Hale, J.S., Boursalian, T.E., Turk, G.L. & Fink, P.J. Thymic output in aged mice. Proc Natl Acad Sci U S A 103, 8447–8452 (2006).

21. Zhang, B. et al. Glimpse of natural selection of long-lived T-cell clones in healthy life. Proc Natl Acad Sci U S A 113, 9858–9863 (2016).

22. Martinet, K.Z., Bloquet, S. & Bourgeois, C. Ageing combines CD4 T cell lymphopenia in secondary lymphoid organs and T cell accumulation in gut associated lymphoid tissue. Immun Ageing 11, 8 (2014).

23. Wertheimer, A.M. et al. Aging and cytomegalovirus infection differentially and jointly affect distinct circulating T cell subsets in humans. J Immunol 192, 2143–2155 (2014).

24. Czesnikiewicz-Guzik, M. et al. T cell subset-specific susceptibility to aging. Clin Immunol 127, 107–118 (2008).

25. Scollay, R.G., Butcher, E.C. & Weissman, I.L. Thymus cell migration. Quantitative aspects of cellular traffic from the thymus to the periphery in mice. Eur J Immunol 10, 210–218 (1980).

26. Yu, W. et al. Continued RAG expression in late stages of B cell development and no apparent re-induction after immunization. Nature 400, 682–687 (1999).

27. Douek, D.C. et al. Changes in thymic function with age and during the treatment of HIV infection. Nature 396, 690–695 (1998).

28. Harrell, M.I., Iritani, B.M. & Ruddell, A. Lymph node mapping in the mouse. J Immunol Methods 332, 170–174 (2008).

29. Tilney, N.L. Patterns of lymphatic drainage in the adult laboratory rat. J Anat 109, 369–383 (1971).

30. Cyster, J.G. Chemokines, sphingosine-1-phosphate, and cell migration in secondary lymphoid organs. Annu Rev Immunol 23, 127–159 (2005).

31. Haluszczak, C. et al. The antigen-specific CD8+ T cell repertoire in unimmunized mice includes memory phenotype cells bearing markers of homeostatic expansion. J Exp Med 206, 435–448 (2009).

32. Quinn, K.M. et al. Age-Related Decline in Primary CD8(+) T Cell Responses Is Associated with the Development of Senescence in Virtual Memory CD8(+) T Cells. Cell Rep 23, 3512–3524 (2018).

33. Renkema, K.R., Li, G., Wu, A., Smithey, M.J. & Nikolich-Žugich, J. Two separate defects affecting true naive or virtual memory T cell precursors combine to reduce naive T cell responses with aging. Journal of immunology (Baltimore, Md.: 1950) 192, 151–159 (2014).

34. Rudd, B.D. et al. Nonrandom attrition of the naive CD8+ T-cell pool with aging governed by T-cell receptor:pMHC interactions. Proc Natl Acad Sci U S A 108, 13694–13699 (2011).

35. Decman, V. et al. Defective CD8 T cell responses in aged mice are due to quantitative and qualitative changes in virus-specific precursors. J Immunol 188, 1933–1941 (2012).

36. Chiu, B.C., Martin, B.E., Stolberg, V.R. & Chensue, S.W. Cutting edge: Central memory CD8 T cells in aged mice are virtual memory cells. J Immunol 191, 5793–5796 (2013).

37. Widman, D.G., Ishikawa, T., Fayzulin, R., Bourne, N. & Mason, P.W. Construction and characterization of a second-generation pseudoinfectious West Nile virus vaccine propagated using a new cultivation system. Vaccine 26, 2762–2771 (2008).

38. Uhrlaub, J.L., Brien, J.D., Widman, D.G., Mason, P.W. & Nikolich-Zugich, J. Repeated in vivo stimulation of T and B cell responses in old mice generates protective immunity against lethal West Nile virus encephalitis. Journal of immunology (Baltimore, Md.: 1950) 186, 3882–3891 (2011).

39. Brien, J.D., Uhrlaub, J.L. & Nikolich-Zugich, J. Protective capacity and epitope specificity of CD8(+) T cells responding to lethal West Nile virus infection. Eur J Immunol 37, 1855–1863 (2007).

40. Linton, P.J. & Dorshkind, K. Age-related changes in lymphocyte development and function. Nat Immunol 5, 133–139 (2004).

41. van Hoeven, V. et al. Dynamics of Recent Thymic Emigrants in Young Adult Mice. Front Immunol 8, 933 (2017).

42. Houston, E.G., Jr., Nechanitzky, R. & Fink, P.J. Cutting edge: Contact with secondary lymphoid organs drives postthymic T cell maturation. J Immunol 181, 5213–5217 (2008).

43. Watanabe, N. et al. Human thymic stromal lymphopoietin promotes dendritic cell-mediated CD4+ T cell homeostatic expansion. Nat Immunol 5, 426–434 (2004).

44. Braun, A. et al. Afferent lymph-derived T cells and DCs use different chemokine receptor CCR7-dependent routes for entry into the lymph node and intranodal migration. Nat Immunol 12, 879–887 (2011).

45. Bromley, S.K., Thomas, S.Y. & Luster, A.D. Chemokine receptor CCR7 guides T cell exit from peripheral tissues and entry into afferent lymphatics. Nat Immunol 6, 895–901 (2005).

46. Pham, T.H., Okada, T., Matloubian, M., Lo, C.G. & Cyster, J.G. S1P1 receptor signaling overrides retention mediated by G alpha i-coupled receptors to promote T cell egress. Immunity 28, 122–133 (2008).

47. Davies, J.S., Thompson, H.L., Pulko, V., Padilla Torres, J. & Nikolich-Zugich, J. Role of Cell-Intrinsic and Environmental Age-Related Changes in Altered Maintenance of Murine T Cells in Lymphoid Organs. J Gerontol A Biol Sci Med Sci 73, 1018–1026 (2018).

48. Alpdogan, S.O. et al. Rapidly proliferating CD44hi peripheral T cells undergo apoptosis and delay posttransplantation T-cell reconstitution after allogeneic bone marrow transplantation. Blood 112, 4755–4764 (2008).

49. Fernandes, S.E., Alakesh, A., Rajmani, R.S., Jhunjhunwala, S. & Saini, D.K. Aging associated altered response to intracellular bacterial infections and its implication on the host. Biochim Biophys Acta Mol Cell Res 1868, 119063 (2021).

50. Smithey, M.J. et al. Lost in translation: mice, men and cutaneous immunity in old age. Biogerontology 16, 203–208 (2015).

51. Fletcher, A.L. et al. Reproducible isolation of lymph node stromal cells reveals site-dependent differences in fibroblastic reticular cells. Front Immunol 2, 35 (2011).

52. Smithey, M.J., Renkema, K.R., Rudd, B.D. & Nikolich-Zugich, J. Increased apoptosis, curtailed expansion and incomplete differentiation of CD8+ T cells combine to decrease clearance of L. monocytogenes in old mice. Eur J Immunol 41, 1352–1364 (2011).

