## Supplementary information for "Early age-related atrophy of cutaneous lymph nodes precipitates an early functional decline in skin immunity in mice with aging"

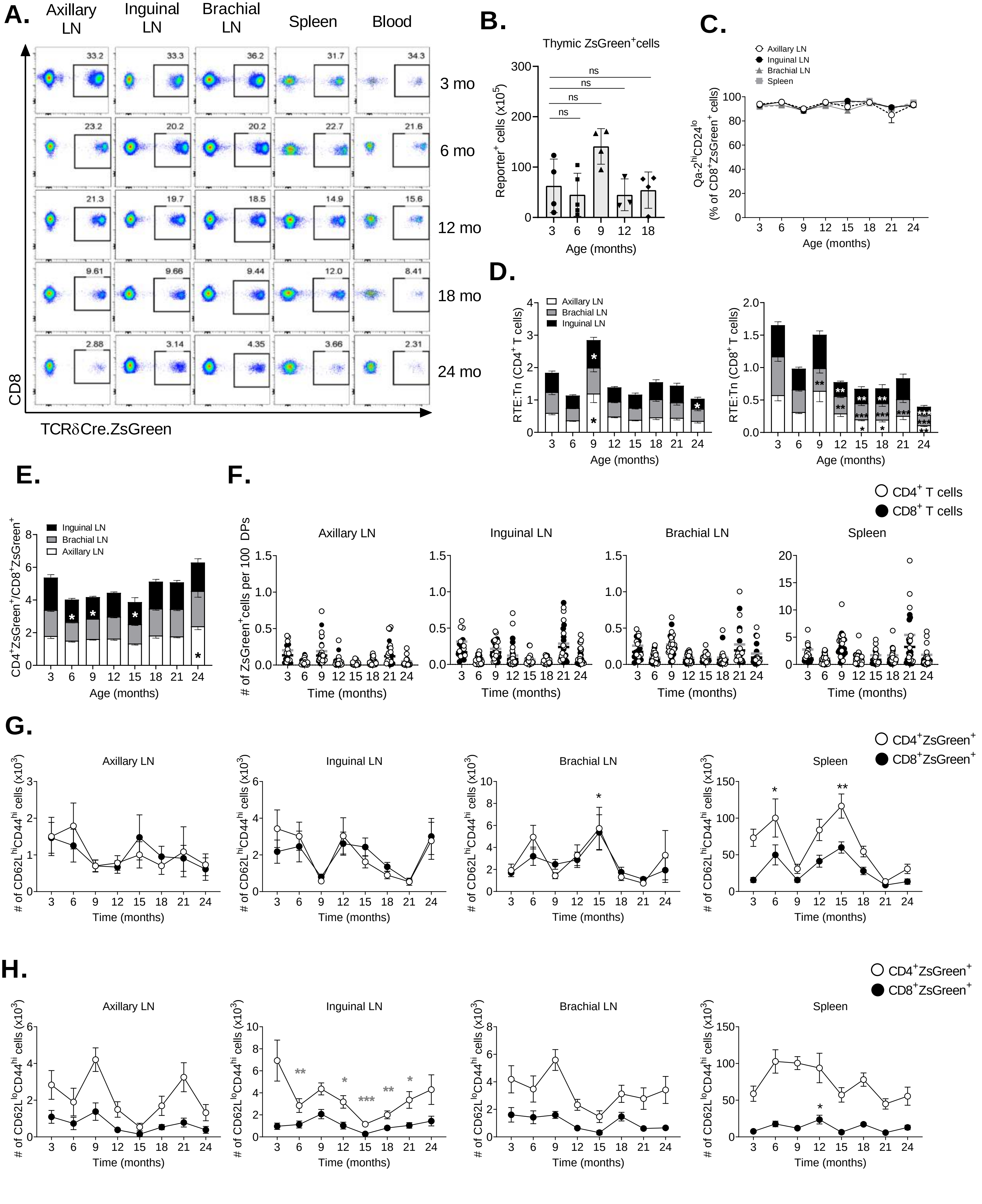
A.
Axillary LN
Inguinal LN
Brachial LN
Spleen
Blood
3 mo
6 mo
12 mo
18 mo
24 mo
CD8
TCRδCre.ZsGreen
B.
Thymic ZsGreen+cells
Reporter+ cells (x10^5)
Age (months)
ns
C.
Axillary LN
Inguinal LN
Brachial LN
Spleen
Qa-2hiCD24lo (% of CD8+ZsGreen+ cells)
Age (months)
D.
RTE:Tn (CD4+ T cells)
RTE:Tn (CD8+ T cells)
Age (months)
E.
Inguinal LN
Brachial LN
Axillary LN
CD4+ZsGreen+/CD8+ZsGreen+
Age (months)
F.
CD4+ T cells
CD8+ T cells
Axillary LN
Inguinal LN
Brachial LN
Spleen
### of ZsGreen+cells per 100 DPs
Time (months)
G.
CD4+ZsGreen+
CD8+ZsGreen+
### of CD62LhiCD44hi cells (x10^3)
Time (months)
H.
### of CD62LloCD44hi cells (x10^3)
Time (months)

**Figure S1**

**Figure S1. Timeline of seeding and retention of newly generated recent thymic emigrants into the SLO.** TCRδ$^{CreER}$.ZsGreen mice were treated as indicated in figure 1A and ZsGreen$^{+}$ RTE were analyzed using multicolor FCM. **(A)** Representative FCM dot plot show ZsGreen-expressing CD8$^{+}$ T cells in different SLO at indicated ages. The numbers within the plot show percentages of indicated cell populations. **(B)** Absolute numbers of ZsGreen$^{+}$ T cells in the thymus at indicated ages are shown. **(C)** Data represent percentages of mature phenotype (Qa2$^{hi}$CD24$^{lo}$) RTE among the CD8$^{+}$ZsGreen$^{+}$ cells of SLO at the indicated ages. **(D)** RTE (ZsGreen$^{+}$) to $T_N$ (CD62L$^{hi}$CD44$^{lo}$) cell ratio among CD4$^{+}$ T cells (left) and CD8$^{+}$ T cells (right) were calculated and plotted. **(E)** Data represent the ratio of absolute numbers of CD4$^{+}$ZsGreen$^{+}$ and CD8$^{+}$ZsGreen$^{+}$ cells of the indicated SLO. **(F)** Absolute numbers of CD4$^{+}$ZsGreen$^{+}$ and CD8$^{+}$ZsGreen$^{+}$ cells normalized to per 100 CD4$^{+}$CD8$^{+}$ double positive (DP) thymocytes from the same mice were shown. Horizontal lines indicate mean of the group (light grey line, CD4$^{+}$ T cells; black line, CD8$^{+}$ T cells). Data show absolute numbers of **(G)** ZsGreen$^{+}$CD62L$^{hi}$CD44$^{hi}$ $T_{CM}$ phenotype and **(H)** ZsGreen$^{+}$CD62L$^{lo}$CD44$^{hi}$ $T_{EM}$ phenotype RTE among the CD4$^{+}$ and CD8$^{+}$ compartments of SLO at the indicated ages. Each dot represents individual mouse **(B, F)** and data indicate mean±SEM of pooled results of three independent experiments with 9-19 mice/age group **(C-H)**. * $p < 0.05$, ** $p < 0.01$, *** $p < 0.001$, **** $p \leq 0.0001$ (p-values for CD4$^{+}$ and CD8$^{+}$ T cells are denoted by grey and black stars, respectively); One-way ANOVA followed by Dunnett's multiple comparison test (compared to 3 mo) **(B-E)**. One-way ANOVA followed by Dunnett's multiple comparison test (compared to 3 mo) **(F-H)**. All the groups were compared to the 3 mo.

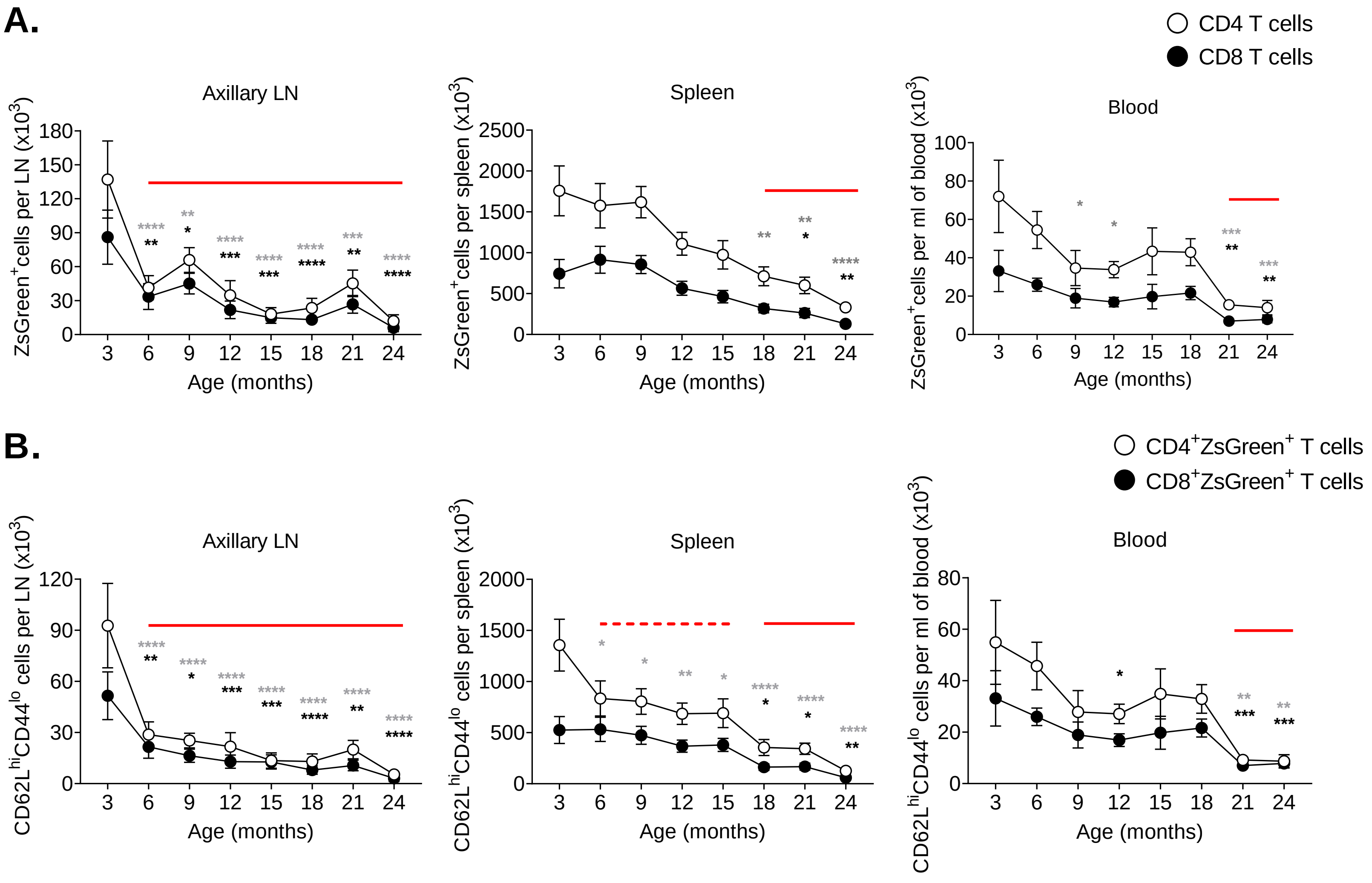
A.
CD4 T cells
CD8 T cells
Axillary LN
Spleen
Blood
ZsGreen+cells per LN (x10³)
ZsGreen+cells per spleen (x10³)
ZsGreen+cells per ml of blood (x10³)
Age (months)
B.
CD4+ZsGreen+ T cells
CD8+ZsGreen+ T cells
CD62LhiCD44lo cells per LN (x10³)
CD62LhiCD44lo cells per spleen (x10³)
CD62LhiCD44lo cells per ml of blood (x10³)

### Figure S2

**Figure S2. Differential kinetics of seeding and retention of newly generated recent thymic emigrants into the SLO.** The SLO of mice from Figure 1A, were analyzed for the cell phenotyping using multicolor FCM. Data show absolute numbers of **(A)** $ZsGreen^{+}$ and **(B)** $ZsGreen^{+}CD62L^{hi}CD44^{lo}$ naive phenotype RTE among the $CD4^{+}$ and $CD8^{+}$ compartments of the indicated tissue. Data represent pooled results of cross-sectional experiment performed across 5-6 independent harvests with 9-19 mice/age group **(A, B)**. Error bar represents mean ± SEM. * $p < 0.05$, ** $p < 0.01$, *** $p < 0.001$, **** $p \leq 0.0001$ (p-values for $CD4^{+}$ and $CD8^{+}$ T cells are denoted by grey and black stars, respectively); Two-way ANOVA followed by Dunnett's multiple comparison test (compared to 3 mo). **(A, B)**. The horizontal red line in the graph denotes the time where significant drop in the number of RTE observed.

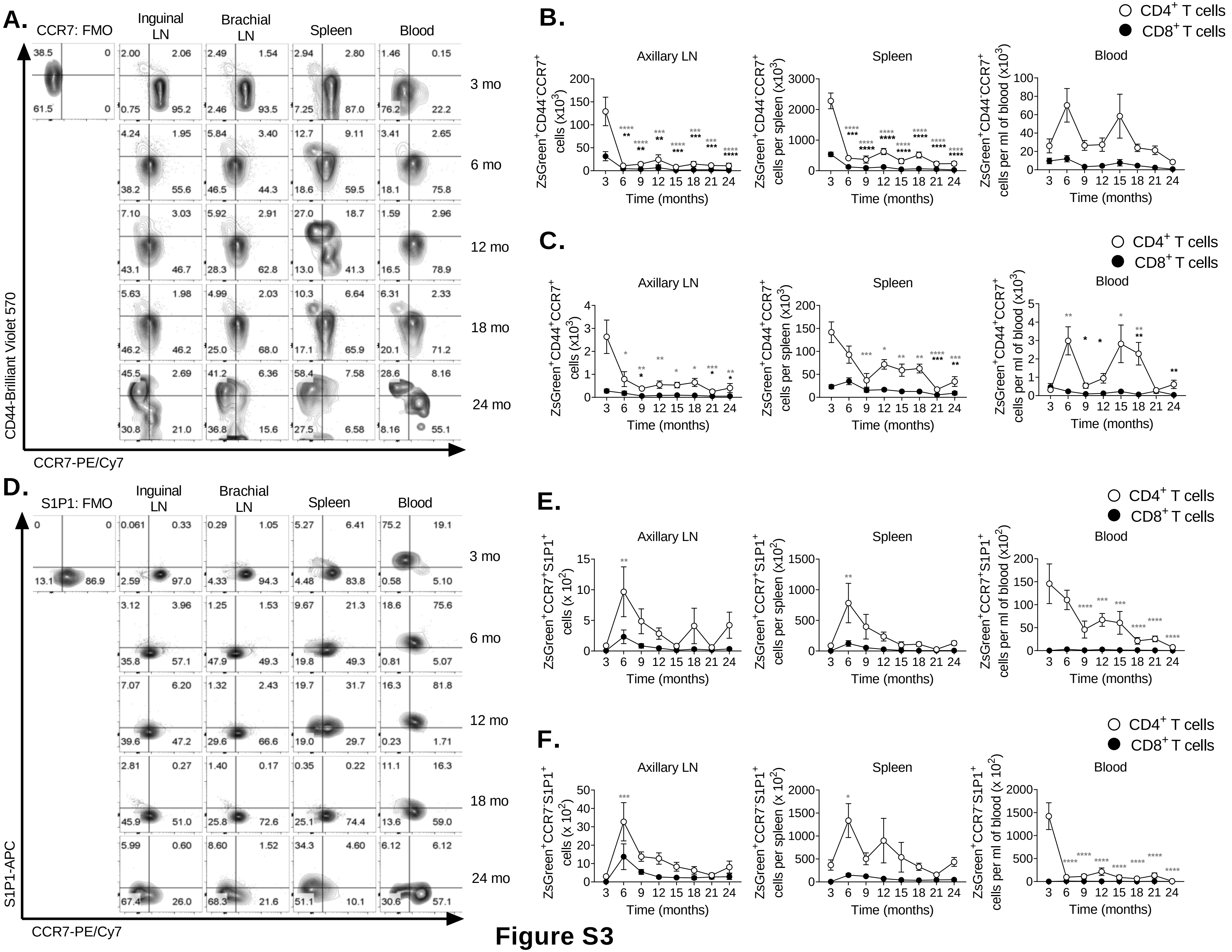

A.
CCR7: FMO
Inguinal LN
Brachial LN
Spleen
Blood
3 mo
6 mo
12 mo
18 mo
24 mo
CD44-Brilliant Violet 570
CCR7-PE/Cy7
B.
C.
CD4+ T cells
CD8+ T cells
Axillary LN
Spleen
Blood
ZsGreen+CD44-CCR7+ cells (x10^3)
ZsGreen+CD44-CCR7+ cells per spleen (x10^3)
ZsGreen+CD44-CCR7+ cells per ml of blood (x10^3)
ZsGreen+CD44+CCR7+ cells (x10^3)
ZsGreen+CD44+CCR7+ cells per spleen (x10^3)
ZsGreen+CD44+CCR7+ cells per ml of blood (x10^3)
Time (months)
D.
S1P1: FMO
S1P1-APC
E.
F.
ZsGreen+CCR7+S1P1+ cells (x 10^2)
ZsGreen+CCR7+S1P1+ cells per spleen (x10^2)
ZsGreen+CCR7+S1P1+ cells per ml of blood (x10^2)
ZsGreen+CCR7-S1P1+ cells (x 10^2)
ZsGreen+CCR7-S1P1+ cells per spleen (x 10^2)
ZsGreen+CCR7-S1P1+ cells per ml of blood (x 10^2)

**Figure S3**

**Figure S3. Decline of CCR7-expressing cells and transient increases in S1P1+ RTE correlates with age-related decline of RTE in SLO.** TCR$\delta^{CreER}$.ZsGreen mice were treated as in figure 1A. Peripheral LN, spleen and blood cells were stained with CCR7 and S1P1 and analyzed by FCM. **(A)** Representative FACS plot show staining of CCR7 and CD44 on cells gated on live, singlet, $CD8^+$, $ZsGreen^+$ T cells. **(B, C)** Data show absolute numbers of **(B)** $ZsGreen^+CD44^-CCR7^+$ cells and **(C)** $ZsGreen^+CD44^+CCR7^+$ cells from CD4 (open circle) and CD8 (closed circle) T cell compartment in the inguinal and brachial LN. **(D)** Representative FACS plot show staining of CCR7 and S1P1 on cells gated on live, singlet, $CD8^+$, $ZsGreen^+$ T cells. **(E, F)** Absolute numbers of **(E)** $ZsGreen^+CCR7^+S1P1^+$ cells and **(F)** $ZsGreen^+CCR7^-S1P1^+$ cells from CD4 (open circle) and CD8 (closed circle) T cell compartment of the SLO and circulation were shown. Data represent pooled results of longitudinal experiment performed across 5-6 independent harvests with 9-19 mice/age group **(B-C, E-F)**. Error bars represent mean±SEM. * $p < 0.05$, ** $p < 0.01$, *** $p < 0.001$, **** $p \leq 0.0001$ (p-values for $CD4^+$ and $CD8^+$ T cells are denoted by grey and black stars, respectively); Two-way ANOVA followed by Dunnett's multiple comparison test (multiple comparisons to 3 mo) **(B-C, E-F)**. Numbers in the graph denotes the percentages of the indicated cell populations **(A, D)**.

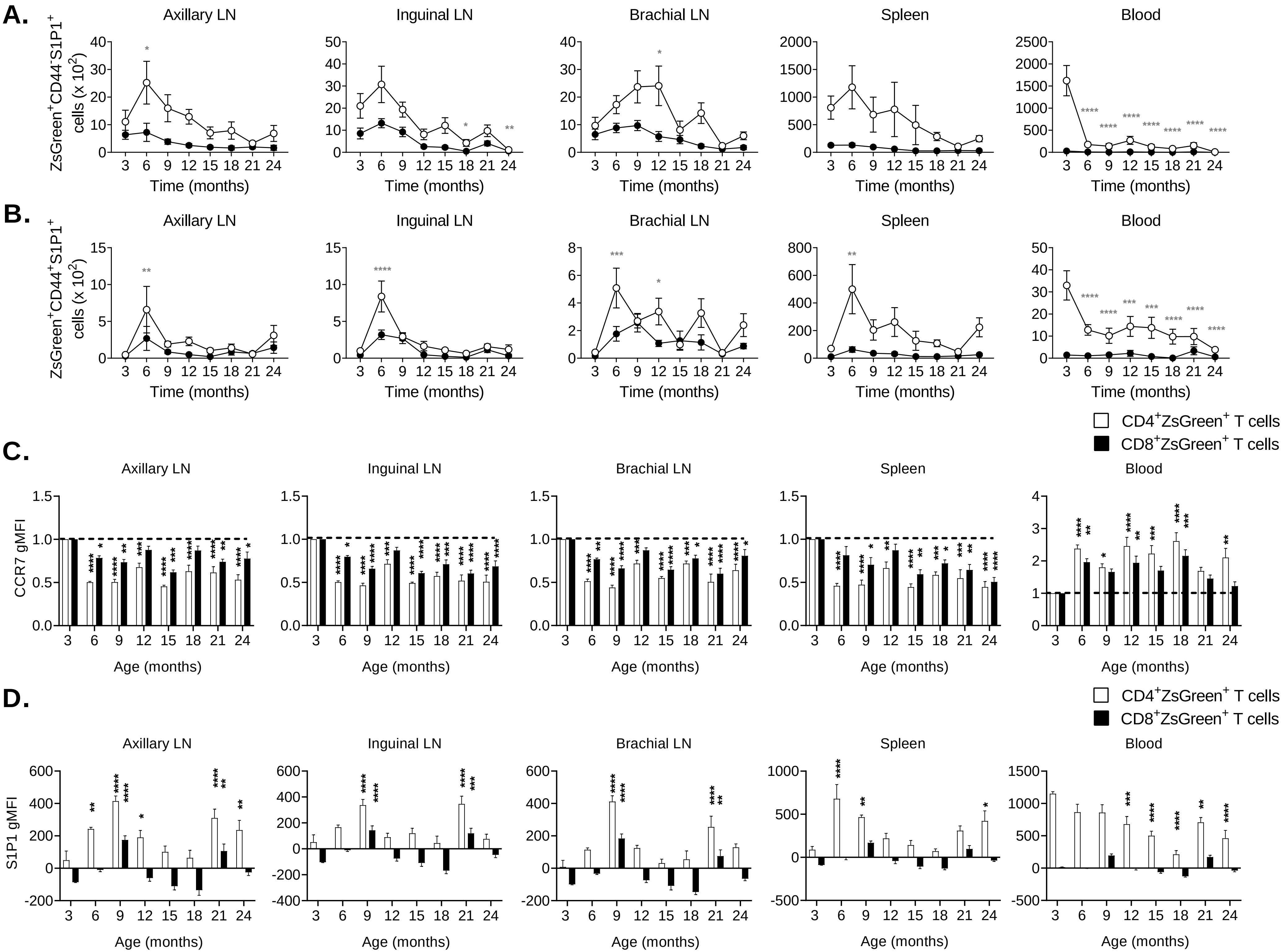
A.
B.
C.
D.
Axillary LN
Inguinal LN
Brachial LN
Spleen
Blood
ZsGreen+CD44-S1P1+ cells (x 10^2)
ZsGreen+CD44+S1P1+ cells (x 10^2)
Time (months)
CCR7 gMFI
S1P1 gMFI
Age (months)
CD4+ZsGreen+ T cells
CD8+ZsGreen+ T cells

**Figure S4**

**Figure S4. Surface expression of CCR7 and S1P1 on RTE in SLO.** TCRδ$^{CreER}$.ZsGreen mice were treated as in figure 1A. Peripheral LN, spleen and blood cells were stained with CCR7 and S1P1 and analyzed by FCM as in figure 2. **(A, B)** Data show absolute numbers of **(A)** ZsGreen$^+$CD44$^-$S1P1$^+$ cells and **(B)** ZsGreen$^+$CD44$^+$S1P1$^+$ cells from CD4 (open circle) and CD8 (closed circle) T cell compartment in the indicated tissue. **(C, D)** The geometric mean fluorescence intensity (gMFI) of **(C)** CCR7 and **(D)** S1P1 on the surface of CD4$^+$ZsGreen$^+$ (open bars) and CD8$^+$ZsGreen$^+$ (black bars) cells were calculated and plotted. The gMFI of CCR7 and S1P1 on cells analyzed on each day were compared to the gMFI of the bead-bound anti-mouse CCR7-PE/Cy7 and anti-mouse S1P1-APC, respectively. The values of gMFI of CCR7 and S1P1 were normalized with the values at 3 mo and plotted. Data represent pooled results of longitudinal experiment performed across 5-6 independent harvests with 9-19 mice/age group **(A-D)**. Error bars represent mean ± SEM. * $p < 0.05$, ** $p < 0.01$, *** $p < 0.001$, **** $p \le 0.0001$; Two-way ANOVA followed by Dunnett's multiple comparison test (compared to 3 mo) **(A-D)**.

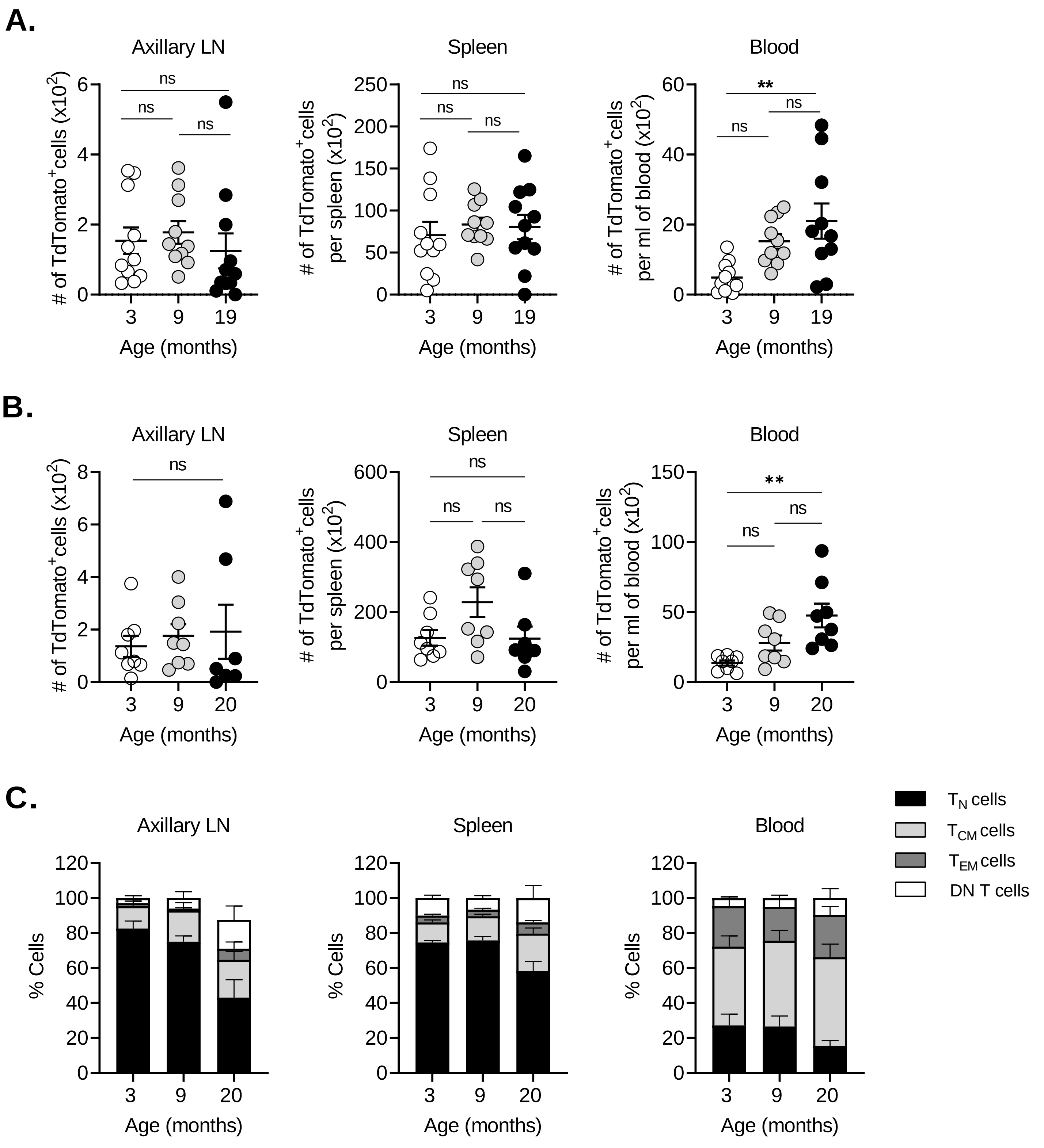
A.
Axillary LN
Spleen
Blood
### of TdTomato+cells (x10^2)
### of TdTomato+cells per spleen (x10^2)
### of TdTomato+cells per ml of blood (x10^2)
ns
**
Age (months)
B.
C.
% Cells
TN cells
TCM cells
TEM cells
DN T cells

**Figure S5**

**Figure S5. Recent thymic emigrants effectively home to the aged peripheral lymph nodes.** TCR$\delta^{CreER}$.TdTomato mice were kept on tamoxifen-containing diet for 30 days followed by 21 days on the normal chow before TdTomato$^{+}$ T cells from the spleen and pLN were FACS sorted at 3 mo of age. About 2 x $10^5$ TdTomato$^{+}$ total T cells were *i.v.* transferred into 3, 9, and 19 months old naïve C57BL/6 mice. **(A)** One hour later and **(B)** 30 days later, trafficking of transferred cells in the SLO were analyzed. Data show absolute numbers of donor TdTomato$^{+}$ RTE in the indicated organ. **(C)** TdTomato$^{+}$ T cells from figure 3B were further analyzed for naïve (CD62L$^{hi}$CD44$^{lo}$), central memory (CD62L$^{hi}$CD44$^{hi}$), effector memory (CD62L$^{lo}$CD44$^{hi}$), and double-negative (DN; CD62L$^{lo}$CD44$^{lo}$) phenotypes and percentages of cell populations among TdTomato$^{+}$ T cells plotted. Data shown are the pool result of two independent transfer experiments with n=8-11 mice per group **(A-C)**. Each dot represents individual mouse **(A-B)**; data represents mean ± SEM **(A-C)**. * $p < 0.05$, ** $p < 0.01$, ns = non-significant. One-way ANOVA followed by Tukey's test **(A, B)**.

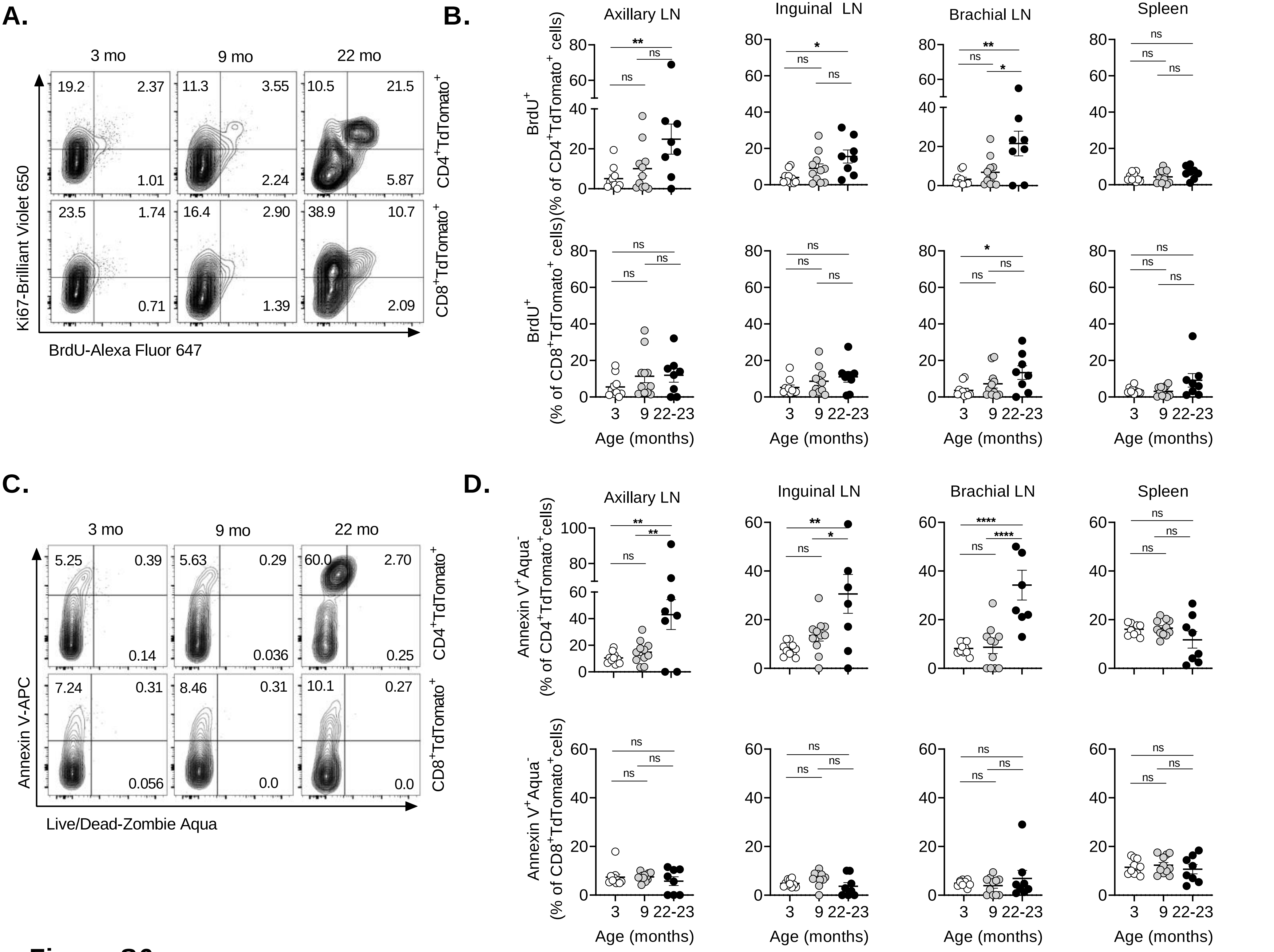
A.
3 mo
9 mo
22 mo
Ki67-Brilliant Violet 650
BrdU-Alexa Fluor 647
CD4+TdTomato+
CD8+TdTomato+
19.2
2.37
1.01
11.3
3.55
2.24
10.5
21.5
5.87
23.5
1.74
0.71
16.4
2.90
1.39
38.9
10.7
2.09
B.
Axillary LN
Inguinal LN
Brachial LN
Spleen
BrdU+ (% of CD4+TdTomato+ cells)
BrdU+ (% of CD8+TdTomato+ cells)
Age (months)
3
9
22-23
C.
Annexin V-APC
Live/Dead-Zombie Aqua
5.25
0.39
0.14
5.63
0.29
0.036
60.0
2.70
0.25
7.24
0.31
0.056
8.46
0.31
0.0
10.1
0.27
0.0
D.
Annexin V+Aqua- (% of CD4+TdTomato+cells)
Annexin V+Aqua- (% of CD8+TdTomato+cells)

**Figure S6**

**Figure S6. Recent thymic emigrants exhibit increased proliferation and apoptosis in old SLO.** TCRδ$^{CreER}$.ZsGreen or TCRδ$^{CreER}$.TdTomato mice were fed with TAM chow for 30 days followed by normal chow and 1 mg/ml 5-bromo-2'-deoxyuridine (BrdU) in drinking water containing 1% glucose for 21 days before mice were sacrificed at 3, 9, and 19 mo. The pLN and spleen cells were intracellularly stained with anti-mouse Ki67 and anti-BrdU, and annexin-V and fixable live/dead-Zombie Aqua. **(A)** Representative FACS plots shows the BrdU and Ki67 stained cells of the inguinal LN. **(B)** BrdU$^{+}$ population frequencies of CD4$^{+}$TdTomato$^{+}$ (top row) and CD8$^{+}$TdTomato$^{+}$ (bottom row) cells of the SLO were shown. **(C)** Representative FACS plots shows cells of the inguinal LN stained with Annexin-V and live/dead-Zombie aqua. **(D)** Annexin-V$^{+}$Zombie Aqua$^{-}$ population frequencies of CD4$^{+}$TdTomato$^{+}$ (top row) and CD8$^{+}$TdTomato$^{+}$ (bottom row) apoptotic cells of the SLO were shown. Numbers in the quadrants represents percentages of the cells **(A, C)**. Each dot represents individual mouse **(B, D)**. Data shown were the pool of two independent experiments with 8-11 mice/group. Error bars represent mean ± SEM. * $p < 0.05$, ** $p < 0.01$, *** $p < 0.001$, **** $p \leq 0.0001$, ns = non-significant; One-way ANOVA followed by Tukey's multiple comparison test **(B, D)**.

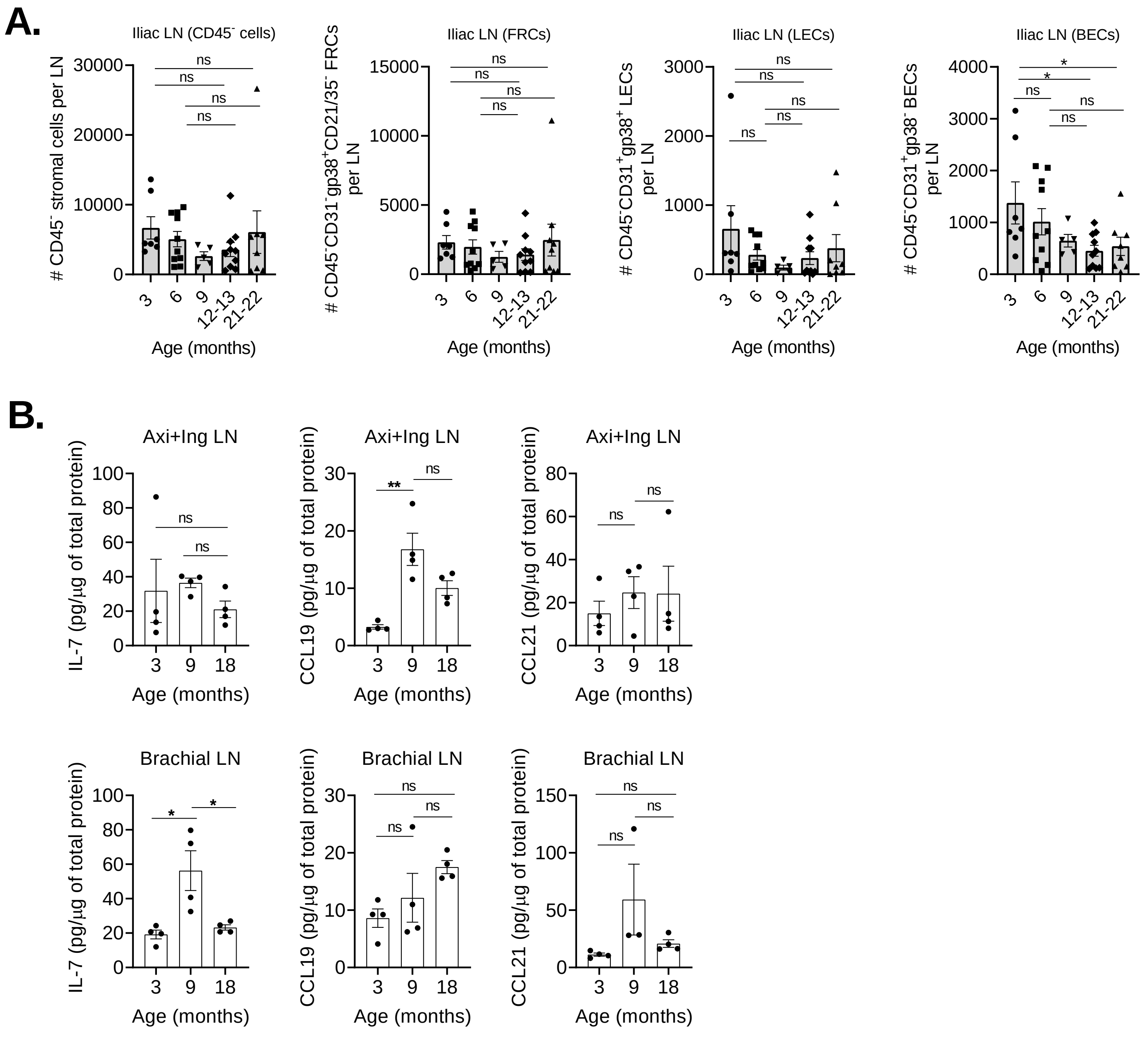
A.
Iliac LN (CD45⁻ cells)
### CD45⁻ stromal cells per LN
Iliac LN (FRCs)
### CD45⁻CD31⁻gp38⁺CD21/35⁻ FRCs per LN
Iliac LN (LECs)
### CD45⁻CD31⁺gp38⁺ LECs per LN
Iliac LN (BECs)
### CD45⁻CD31⁺gp38⁻ BECs per LN
3
6
9
12-13
21-22
Age (months)
ns
*
B.
Axi+Ing LN
Brachial LN
IL-7 (pg/µg of total protein)
CCL19 (pg/µg of total protein)
CCL21 (pg/µg of total protein)
3
9
18
Age (months)
ns
*
**

### Figure S7

**Figure S7. Enumeration of stromal cells and survival or tropic factors in the peripheral lymph nodes during aging. (A)** Pooled Iliac LNs of the naive wild-type C57BL/6 mice of indicated ages were digested with Liberase-TL and DNase-I for stromal cell analysis as in figure 4A and stromal cells were analyzed by FCM. Data show absolute numbers of total stromal cells ($CD45^-$ cells), FRCs ($CD45^-CD31^-gp38^+CD21^-CD35^-$), LECs ($CD45^-CD31^+gp38^+$), and BECs ($CD45^-CD31^+gp38^-$). n= 5-10 mice/age group. **(B)** IL-7, CCL19, and CCL protein levels normalized to total protein content of whole LN lysate were shown. n= 4 mice/group. Each dot represents individual mouse and data indicate mean±SEM. (A). * $p < 0.05$, ** $p < 0.01$; One-way ANOVA followed by Dunnett's multiple comparison test **(A)**, One-way ANOVA followed by Tukey's multiple comparison test **(B)**.

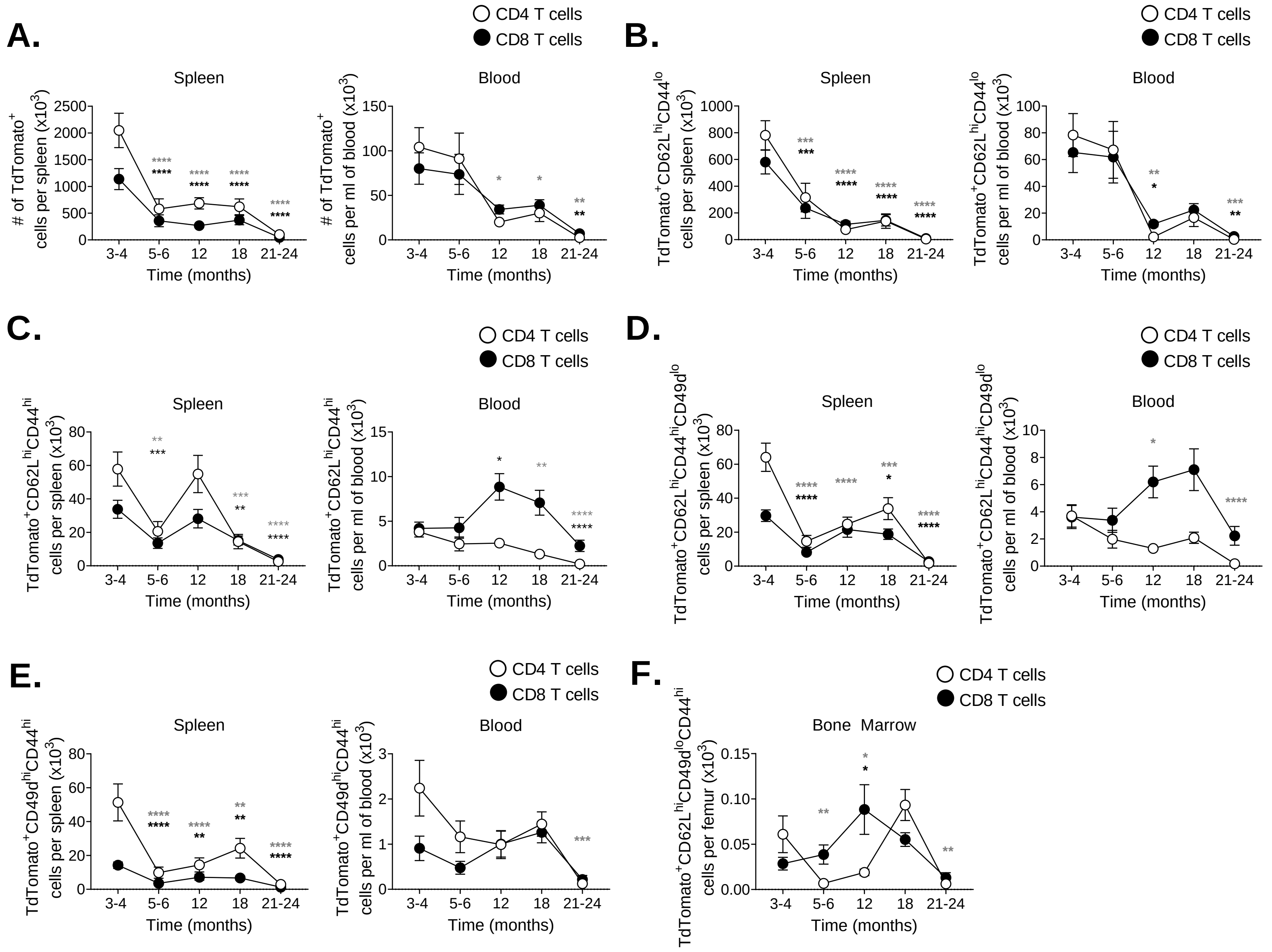
A.
CD4 T cells
CD8 T cells
Spleen
### of TdTomato+ cells per spleen (x10^3)
Time (months)
Blood
### of TdTomato+ cells per ml of blood (x10^3)
B.
Spleen
TdTomato+CD62L^hiCD44^lo cells per spleen (x10^3)
Blood
TdTomato+CD62L^hiCD44^lo cells per ml of blood (x10^3)
C.
TdTomato+CD62L^hiCD44^hi cells per spleen (x10^3)
TdTomato+CD62L^hiCD44^hi cells per ml of blood (x10^3)
D.
TdTomato+CD62L^hiCD44^hiCD49d^lo cells per spleen (x10^3)
TdTomato+CD62L^hiCD44^hiCD49d^lo cells per ml of blood (x10^3)
E.
TdTomato+CD49d^hiCD44^hi cells per spleen (x10^3)
TdTomato+CD49d^hiCD44^hi cells per ml of blood (x10^3)
F.
Bone Marrow
TdTomato+CD62L^hiCD49d^loCD44^hi cells per femur (x10^3)
3-4
5-6
12
18
21-24

### Figure S8

**Figure S8. Age-timeline of time-stamped cells with naïve and memory phenotype in SLO.** TdTomato$^{+}$ time-stamped T cells from the mice in figure 5 were analyzed for naïve and memory phenotype. Data show absolute numbers of **(A)** total TdTomato$^{+}$ time-stamped cells, and **(B)** TdTomato$^{+}$ time-stamped cells with naive (CD62L$^{hi}$CD44$^{lo}$), **(C)** central memory (CD62L$^{hi}$CD44$^{hi}$), **(D)** virtual memory (CD62L$^{hi}$CD44$^{hi}$CD49d$^{lo}$), and **(E)** true memory (CD62L$^{hi}$CD44$^{hi}$CD49d$^{lo}$) phenotype in the spleen and blood are shown. **(F)** Data show absolute numbers of virtual memory cells in the bone marrow. Data represent pooled results of longitudinal experiment performed across 5 independent harvests with 11-18 mice/age group **(A-F)**. Error bar represents mean±SEM. * $p < 0.05$, ** $p < 0.01$, *** $p < 0.001$, **** $p \leq 0.0001$ (p-values for CD4$^{+}$ and CD8$^{+}$ T cells are denoted by grey and black stars, respectively); One-way ANOVA followed by Dunnett's multiple comparison test (compared to 3-4 mo) **(A-F)**.

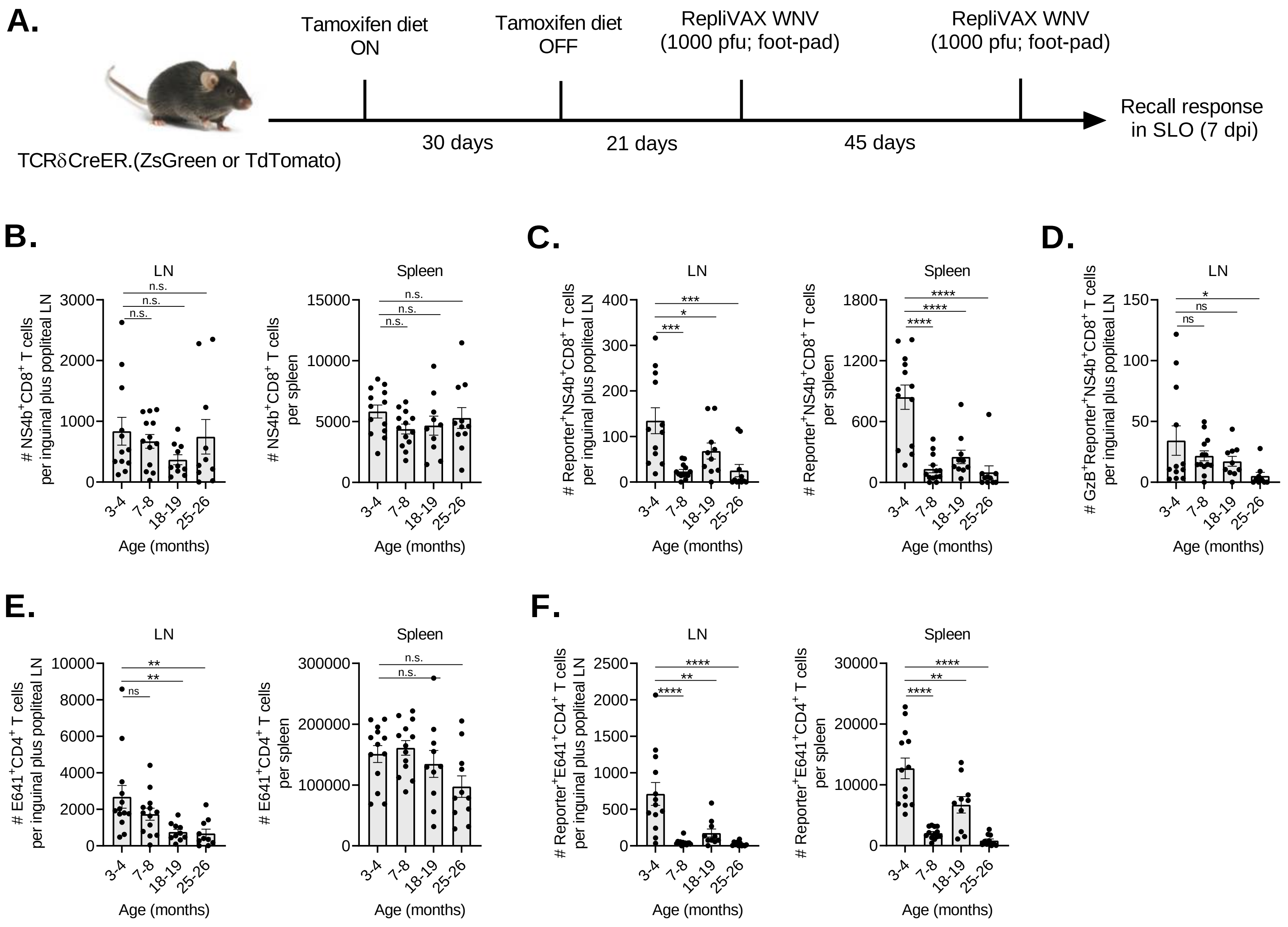
A.
Tamoxifen diet ON
Tamoxifen diet OFF
RepliVAX WNV (1000 pfu; foot-pad)
RepliVAX WNV (1000 pfu; foot-pad)
Recall response in SLO (7 dpi)
TCRδCreER.(ZsGreen or TdTomato)
30 days
21 days
45 days
B.
LN
Spleen
### NS4b+CD8+ T cells per inguinal plus popliteal LN
### NS4b+CD8+ T cells per spleen
n.s.
C.
LN
Spleen
### Reporter+NS4b+CD8+ T cells per inguinal plus popliteal LN
### Reporter+NS4b+CD8+ T cells per spleen
D.
LN
Spleen
### GzB+Reporter+NS4b+CD8+ T cells per inguinal plus popliteal LN
### GzB+Reporter+NS4b+CD8+ T cells per spleen
ns
E.
LN
Spleen
### E641+CD4+ T cells per inguinal plus popliteal LN
### E641+CD4+ T cells per spleen
F.
LN
Spleen
### Reporter+E641+CD4+ T cells per inguinal plus popliteal LN
### Reporter+E641+CD4+ T cells per spleen
3-4
7-8
18-19
25-26
Age (months)

**Figure S9**

**Figure S9. RTE responsive to the Replivax West Nile virus vaccine show detectable recall response. (A)** Pictorial depiction of experimental strategy to assess the recall response of $reporter^{+}$ cells. Briefly, $TCR\delta^{CreER}$.TdTomato mice were fed on tamoxifen-containing diet for 30 days and subsequently maintained on normal diet for next 21 days. At day 21, mice were *s.c.* injected at food-pad with Replivax West Nile virus vaccine (1 x $10^3$ pfu) and received booster dose after 45 days of primary vaccination. At day 7 post booster dose, West Nile virus specific $NS4b^{+}$ and $E641^{+}$ T cell recall response in the $CD8^{+}reporter^{+}$ and $CD4^{+}reporter^{+}$, respectively were analyzed in the SLO. **(B-D)** Data show absolute numbers of **(B)** $NS4b^{+}CD8^{+}$ T cells, **(C)** $reporter^{+}NS4b^{+}CD8^{+}$ T cells, and **(D)** granzyme $B^{+}$reporter+$NS4b^{+}CD8^{+}$ T cells in the pool of draining popliteal and inguinal LN (left) and spleen (right). (E, F) Data show absolute numbers of **(E)** $E641^{+}CD4^{+}$ T cells, **(F)** $reporter^{+}E641^{+}CD4^{+}$ T cells in the pool of draining popliteal and inguinal LN (left) and spleen (right). Data represent pooled results from three separate experiments with 10-13 mice per age group **(B-F)**. Each circle represents individual mice. Error bar represents mean ± SEM. * $p < 0.05$, ** $p < 0.01$, *** $p < 0.001$, **** $p \leq 0.0001$; One-way ANOVA followed by Dunnett's multiple comparison test **(B-F)**.

#### Table S1. Reagents and Resources

| 1. Antibodies | | | |
|---|---|---|---|
| **Molecule** | **Clone** | **Catalogue number** | **Company** |
| Anti-mouse CD3-Alexa fluor 700 | 17A2 | 100215 | Biolegend |
| Anti-mouse CD4-Brilliant Violet 750 | GK1.5 | 100467 | Biolegend |
| Anti-mouse CD4-APC/eF780 | GK1.5 | 47-0042-82 | eBioscience |
| Anti-mouse CD8a-Brilliant Violet 785 | 53-6.7 | 100750 | Biolegend |
| Anti-mouse CD8a-PE | 53-6.7 | 100708 | Biolegend |
| Anti-mouse CD62L-APC/eF780 | MEL-14 | 47-0621-82 | ThermoFisher |
| Anti-mouse/human CD44-Brilliant Violet 570 | IM7 | 103037 | Biolegend |
| Anti-mouse CD5-Brilliant Violet 650 | 53-7.3 | BDB740444 | ThermoFisher |
| Anti-mouse CD49d-PerCp/Cy5.5 | R1-2 | 103620 | Biolegend |
| Anti-mouse CD122-PE | 5H4 | 105906 | Biolegend |
| Anti-mouse Qa-2-Alexa fluor 647 | 695H1-9-9 | 121708 | Biolegend |
| Anti-CD24-PerCp-eF710 | M1/69 | 46-0242-82 | ThermoFisher |
| Anti-mouse CD69-PE-Dazzle 594 | H1.2F3 | 104535 | Biolegend |
| Anti-mouse CD49d-PE | R1-2 | 103608 | Biolegend |
| Anti-mouse/human CD44-Brilliant Violet 421 | IM7 | 103040 | Biolegend |
| Anti-mouse CD5-Brilliant Violet 650 | 53-7.3 | BDB740444 | ThermoFisher |
| Anti-mouse CCR7-PE/Cy7 | 4B12 | 25-1971-82 | eBioscience |
| Anti-mouse S1P1-APC | 713412 | FAB7089P | R&D Systems |
| Anti-mouse CD31-Alexa fluor 647 | 390 | 102415 | Biolegend |
| Anti-mouse PDPN (gp38)-PE | 8.1.1 | 127408 | Biolegend |
| Anti-mouse CD45-eFluor 450 | 30F-11 | 48-0451-80 | eBioscience |
| Anti-mouse Ter119-eFluor 450 | Ter119 | 116235 | Biolegend |
| Anti-mouse CD21/CD35-PerCp/Cy5.5 | 7-E9 | 123416 | Biolegend |
| Anti-mouse MAdCAM1-Alexa fluor 488 | MECA-367 | 120708 | Biolegend |
| Anti-mouse CD3-APC | 17A2 | 100236 | Biolegend |
| Anti-mouse/human Ki-67-Brilliant Violet 650 | 11F6 | 151215 | Biolegend |
| anti-human/mouse Granzyme B-PE/Cy7 | QA16A02 | 372214 | Biolegend |
| Syrian Hamster Anti-podoplanin-Biotin | 8.1.1 | 14-5381-80 | ThermoFisher Scientific |
| Rabbit polyclonal anti-Lyve1 | | ab14917 | Abcam |
| Goat polyclonal anti-rabbit IgG-Alexa fluor 488 | | ab150077 | Abcam |
| Anti-BrdU-Alexa fluor 647 | | 364108 | Biolegend |
| Annexin V-APC | | 640920 | Biolegend |
| **2. Reagents** | **Source** | **Identifier** | |
| Liberase TL | Sigma | Cat # 5401020001 | |
| DNase-I | Sigma | Cat # DN25-100mg | |
| Alexa Fluor™ 555 Tyramide SuperBoost™ Kit, streptavidin | Thermo Fisher Scientific | Cat # B40933 | |
| Mouse CCL19/MIP-3 beta DuoSet ELISA | R & D Systems | Cat # DY440 | |
| Mouse CCL21/6Ckine DuoSet ELISA | R & D Systems | Cat # DY457 | |
| Mouse IL-7 DuoSet ELISA | R & D Systems | Cat # DY407 | |
| ProLong™ Gold Antifade Mountant | Thermo Fisher Scientific | Cat # P10144 | |
| CountBright™ Absolute Counting Beads, for flow cytometry | Thermo Fisher Scientific | Cat # C36950 | |
| CytoFix/CytoPerm kit | BD Biosciences | Cat # 554714 | |
| Pan T cell isolation kit, mouse | Miltenyi Biotec | Cat # 130-095-130 | |
| Tamoxifen diet | Envigo | Cat # TD.1308603 | |
| Zombie Aqua fixable viability kit | Biolegend | Cat # 423102 | |
